# Development and Characterization of a Fixed Repertoire of Blood Transcriptome Modules Based on Co-expression Patterns Across Immunological States

**DOI:** 10.1101/525709

**Authors:** Matthew C Altman, Darawan Rinchai, Nicole Baldwin, Mohammed Toufiq, Elizabeth Whalen, Mathieu Garand, Basirudeen Ahamed Kabeer, Mohamed Alfaki, Scott Presnell, Prasong Khaenam, Aaron Ayllon Benitez, Fleur Mougin, Patricia Thébault, Laurent Chiche, Noemie Jourde-Chiche, J Theodore Phillips, Goran Klintmalm, Anne O’Garra, Matthew Berry, Chloe Bloom, Robert J Wilkinson, Christine M Graham, Marc Lipman, Ganjana Lertmemongkolchai, Davide Bedognetti, Rodolphe Thiebaut, Farrah Kheradmand, Asuncion Mejias, Octavio Ramilo, Karolina Palucka, Virginia Pascual, Jacques Banchereau, Damien Chaussabel

## Abstract

As the capacity for generating large scale data continues to grow the ability to extract meaningful biological knowledge from it remains a limitation. Here we describe the development of a new fixed repertoire of transcriptional modules. It is meant to serve as a stable reusable framework for the analysis and interpretation of blood transcriptome profiling data. It is supported by customized resources, which include analysis workflows, fingerprint grid plots data visualizations, interactive web applications providing access to a vast number of module-specific functional profiling reports, reference transcriptional profiles and give users the ability to visualize of changes in transcript abundance across the modular repertoire at different granularity levels. A use case focusing on a set of six modules comprising interferon-inducible genes is also provided. Altogether we hope that this resource will also serve as a framework for improving over time our collective understanding of the immunobiology underlying blood transcriptome profiling data.

## INTRODUCTION

Technology advancements over the past two decades paired with improvements in cost-effectiveness have greatly increased the capacity to conduct large-scale molecular profiling in biomedical research. In translational settings, these advances have enabled the measurement of molecular phenotypes at very high resolutions, reaching the point where it has become possible to routinely generate whole genome, proteome, metabolome, microbiome and transcriptome profiles on an almost routine basis (1). The current bottleneck is in interpreting the biological significance or potential clinical implications of these molecular signatures.

Our group has developed co-expression modules and used them as a framework to analyze and interpret blood transcriptome data (2–4). Importantly, we have used the modules as a fixed framework: transcriptional modules are not generated each time a new dataset needs to be analyzed; instead, the same set of pre-existing modules is reused to analyze each new dataset. Consequently, our team has released only two different module repertoires over a 12-year period, which we and others have used to analyze numerous blood transcriptome datasets [e.g. (5–8)]. Other blood transcriptome frameworks have been generated (9), as well as those with a more generic transcriptional module repertoire (10). An inherent benefit of using a collection of gene sets is that it permits the reduction of high-dimensional data. Using a fixed repertoire of modules based on co-expression analyses across a large dataset collection, improves the robustness of analyses of small datasets (10). As such, efforts can rather focus on the functional interpretation of the collection of gene sets constituting the repertoire. Furthermore, using the same fixed framework to analyze multiple independent datasets means that cross-study comparisons and interpretation are easier and more reliable. With the construction of this new repertoire we aimed first at increasing the range of immunological states upon which the repertoire is based, which was limited to 7 in our earlier attempt. Secondly, considerable efforts were deployed to improve resources to support analysis and interpretation.

Thus, the range of disease and physiological states employed for the construction of this new modular repertoire has been expended considerably compared to the first and second generation repertoires. Specifically, 16 input datasets were employed, comprising 985 unique blood transcriptome profiles from: patients with autoimmune, infectious or inflammatory diseases; cancer patients; liver transplant recipients; and pregnant mothers. We describe here how we developed the analysis workflows for group comparison and individual molecular “fingerprinting”. Ad hoc data visualization strategies are presented as well, namely the use of module fingerprint grids and heatmaps to represent the data. We also provide access to extensive functional annotations for each of the 382 modules comprising the repertoire via an interactive web application. Reference blood transcriptome fingerprints are also made available via a web application that enables users to explore the changes observed across diseases/studies and among individuals. A focused effort aiming at the characterization of a set of six interferon modules is presented to illustrate the manner in which such resources may be deployed to help unravel the biology underlying blood transcriptome signatures.

## RESULTS

### Constitution of a collection of datasets covering a wide range of immune states

The development of transcriptional module repertoires relies on identifying gene co-expression events using transcriptome profiling data as a starting point. For this new blood transcriptome module repertoire, we used 16 datasets (GEO ID: GSE100150) that encompassed 985 individual whole blood transcriptome profiles. Each dataset corresponds to a different pathological or physiological state (**Table 1**). These datasets were processed in the same facility and run on Illumina HT12 BeadArrays (details are provided in the method section). Similar to our first two repertoires (**Table 2**), we included data from patients (adult and pediatric) with: systemic lupus erythematosus (SLE), systemic onset juvenile idiopathic arthritis (SoJIA), liver transplants and receiving maintenance immunosuppressive therapy, metastatic melanoma, and infectious diseases [with an expanded range that now includes infections caused by influenza, respiratory syncytial virus (RSV), human immunodeficiency virus (HIV) infections, *Mycobacterium tuberculosis, Staphylococcus aureus*, and *Burkholderia pseudomallei (the agent of Melioidosis) and sepsis caused by other bacteria (Streptococcus pneumoniae, Salmonella spp, Pseudomonas aeruginosa)*]. We also added six new conditions to our framework: inflammatory conditions of the skin (juvenile dermatomyositis), lung [chronic obstructive pulmonary disease (COPD)] and circulation (Kawasaki disease); multiple sclerosis (MS); primary immune (B-cell) deficiency; and pregnancy (serving as a physiological variant). Absent from this repertoire (but represented earlier) are patients with type 1 diabetes or with an *Escherichia coli* infection.

**Table 1:** Datasets used for module construction. Sixteen distinct datasets were used as the input for module repertoire construction. Each dataset corresponds to a different condition or physiological state and comprises both cases and matched controls. Each dataset was processed as a single batch at the same facility with the data generated using Illumina HumanHT-12 v3.0 Gene Expression BeadChips. The collection comprises a total of 985 individual transcriptome profiles.

| Dataset | Category | Population | # Samples (Cases) | # Samples (Control) | # Samples (Total) |
| --- | --- | --- | --- | --- | --- |
| 1 *Staphylococcus aureus* infection | Bacterial Infection | Pediatric | 99 | 44 | 143 |
| 2 Sepsis | Bacterial Infection | Adult | 35 | 12 | 47 |
| 3 Tuberculosis | Bacterial Infection | Adult | 23 | 11 | 34 |
| 4 Influenza | Viral Infection | Pediatric | 25 | 14 | 39 |
| 5 RSV | Viral Infection | Pediatric | 70 | 14 | 84 |
| 6 HIV | Viral Infection | Adult | 28 | 35 | 63 |
| 7 Systemic Lupus Erythematosus | Autoimmune | Pediatric | 55 | 14 | 69 |
| 8 Multiple Sclerosis | Autoimmune | Adult | 34 | 22 | 56 |
| 9 Juvenile Dermatomyositis | Autoimmune | Pediatric | 40 | 9 | 49 |
| 10 Kawasaki disease | Autoinflammatory | Pediatric | 21 | 23 | 44 |
| 11 Systemic Onset Idiopathic Arthritis | Autoinflammatory | Pediatric | 62 | 23 | 85 |
| 12 COPD | Inflammatory | Adult | 19 | 24 | 43 |
| 13 Melanoma | Malignancy | Adult | 22 | 5 | 27 |
| 14 Pregnancy | Physiologic variant | Adult | 25 | 20 | 45 |
| 15 Liver transplant recipients | Immunosuppressed | Adult | 94 | 30 | 124 |
| 16 B-cell deficiency | Immunodeficiency | Adult | 20 | 13 | 33 |

**Table 2:** The characteristics of the input datasets used to construct three consecutive generations of blood transcriptome module repertoires.

|  | Generation 1 | Generation 2 | Generation 3 |
| --- | --- | --- | --- |
| **Number of States** | 8 | 7 | 16 |
| **States** |  |  |  |
| Systemic onset juvenile idiopathic arthritis | X | X | X |
| Systemic lupus erythematosus | X | X | X |
| Juvenile dermatomyositis |  |  | X |
| Type 1 Diabetes | X |  |  |
| Multiple sclerosis |  |  | X |
| Kawasaki disease |  |  | X |
| COPD |  |  | X |
| Tuberculosis |  | X | X |
| Sepsis |  | X | X |
| Respiratory Syncytial Virus infection |  |  | X |
| Influenza virus infection | X |  | X |
| Human Immunodeficiency Virus infection |  | X | X |
| *Escherichia coli* infection | X |  |  |
| *Staphylococcus aureus* infection | X |  | X |
| B-cell deficiency |  |  | X |
| Liver transplant recipients | X | X | X |
| Metastatic melanoma | X | X | X |
| Pregnancy |  |  | X |
| **Number of input datasets** | 8 | 9 | 16 |
| **Number of individual profiles** | 239 | 410 | 985 |
| **Sample source** | PBMCs | Whole Blood | Whole Blood |
| **Platform** | Affymetrix U133A&B | Illumina Hu6 v2 | Illumina HT12 v3.0 |
| **Rounds of module selection** | 3 | 8 | 15 |
| **Number of modules** | 28 | 260 | 382 |
| **Year published & reference** | 2008 (4) | 2013 (3) | Current work |

In summary, the sixteen datasets that have been assembled capture a wide breadth of immunological responses. This should permit the construction of a transcriptional module repertoire that will prove useful as a generic framework for blood transcriptome data analyses.

### Implementation of a step-wise approach to blood transcriptional module repertoire construction

After collating the 16 input datasets, we next followed a stepwise process to construct a module repertoire and identify co-expression networks (**Figure 1**). We used the module construction algorithm that we have previously implemented (the code is provided in **Supplementary File 1**). This approach followed four main steps: 1) clustering for each individual dataset; 2) co-clustering, where the number of instances that two genes were included in the same cluster was recorded, with the values ranging from 0 to 16 (i.e. co-clustering observed in none or all of the datasets, respectively); 3) constructing a weighted co-expression network, where the edges between the genes represent at least one co-clustering event in one of the input datasets and the weight is assigned based on the total number of co-clustering events; (**supplementary Figure 1**); and 4) mining the resulting network to identify highly inter-connected sub-networks that form the modules.

**Figure 1:**
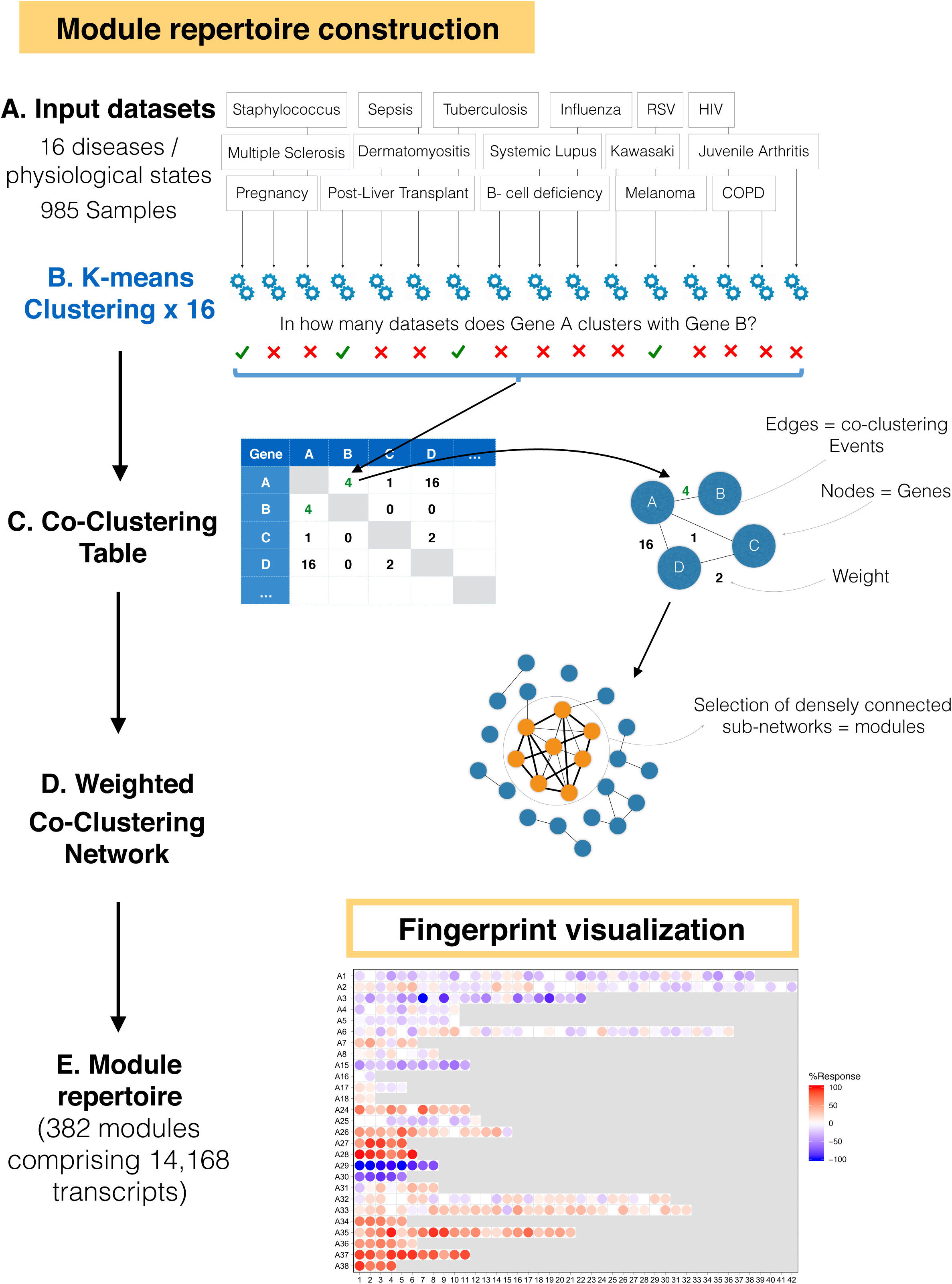
The module repertoire construction process. A. A collection of 16 transcriptome datasets spanning a wide range of immunological and physiological states were used as input. B. Each dataset was independently clustered via k-means clustering. C. Gene co-clustering events were recorded in a table, where the entries indicate the number of datasets in which co-clustering was observed for a given gene pair. D. The co-clustering table served as the input to a weighted co-clustering graph (see also **Supplementary Figure 1**), where the nodes represent genes and the edges represent co-clustering events. E. The largest, most highly weighted sub-networks among a large network (here constituting 15,132 nodes) were identified mathematically and assigned a module ID. The genes constituting this module were removed from the selection pool and the process was repeated, resulting in the selection of 382 modules constituted by 14,168 transcripts.

Constituting a module repertoire in this manner is thus entirely data-driven: it does not rely on any *a priori* information about gene interactions or functions. In total, we identified 382 modules comprising 14,168 transcripts (95.8% of the transcripts detected in this dataset collection).

### Development of module-level analysis workflows and visualizations

A key characteristic of the gene sets collected via the process described above is that, by construction, changes in abundance of the corresponding transcripts within a given module will tend to be coordinated. As such, it should be possible to use these modules as a “framework” to: 1) identify functional convergences among the genes that comprise each set, and 2) summarize changes in overall transcript abundance related to pathological processes or therapeutic interventions.

We determined the gene composition of each of the 382 modules (**supplementary file 2)**, The average number of genes per module as 37.1, the median as 26.5 and the range as 12-169. Functional profiling and enrichment results were generated using multiple tools (GSAn, Literature Lab, IPA, DAVID, KEGG, Biocarta, OMIM, and GOTERM). We also determined the extent of overlap with the previous modular repertoire and a set of modules constituted at Emory University and our previous generation of modules (9). For module-level analyses, we determined the proportion of the constitutive transcripts that differ in abundance levels between study groups (e.g. cases vs. controls; pre-treatment vs. post-treatment). By this approach, two values, corresponding to the percent of transcripts that are (i) increased and (ii) decreased, are derived. The cutoff points can be chosen based on user preferences. For example, cutoffs can be based on statistics, fold changes and/or differences with or without multiple testing correction for group comparisons.

From here, the extent of differential expression at the module level can be displayed as a “fingerprint”, assigning each module to a fixed position on a grid plot and color-coding it according to the level of increased or decreased abundance for the constituting transcripts (**Figure 2**). For this we then performed a second tier of clustering to group the 382 modules into 38 “aggregates”, with each row on the grid displaying the modules corresponding to one such aggregate. Segregation into distinct aggregates was based on similarities in abundance levels observed across the collection of 16 datasets. By this approach, we derived two levels of granularity (i.e. module-level vs. module aggregate-level) with the number of variables for interpretation constrained to a more manageable number. The overall result is that changes in expression levels for each row on the fingerprint grid will tend to be coordinated, which was not the case of prior iterations of such fingerprint grids (**Figure 3**). Some degree of functional convergence can thus be observed within a given row of modules. As an example, in our fingerprint we found that row A1 comprised several modules associated with lymphocytes, while row A28 comprised six distinct “interferon modules” and rows A33 and A35 comprised a number of modules functionally associated with inflammation.

**Figure 2:**
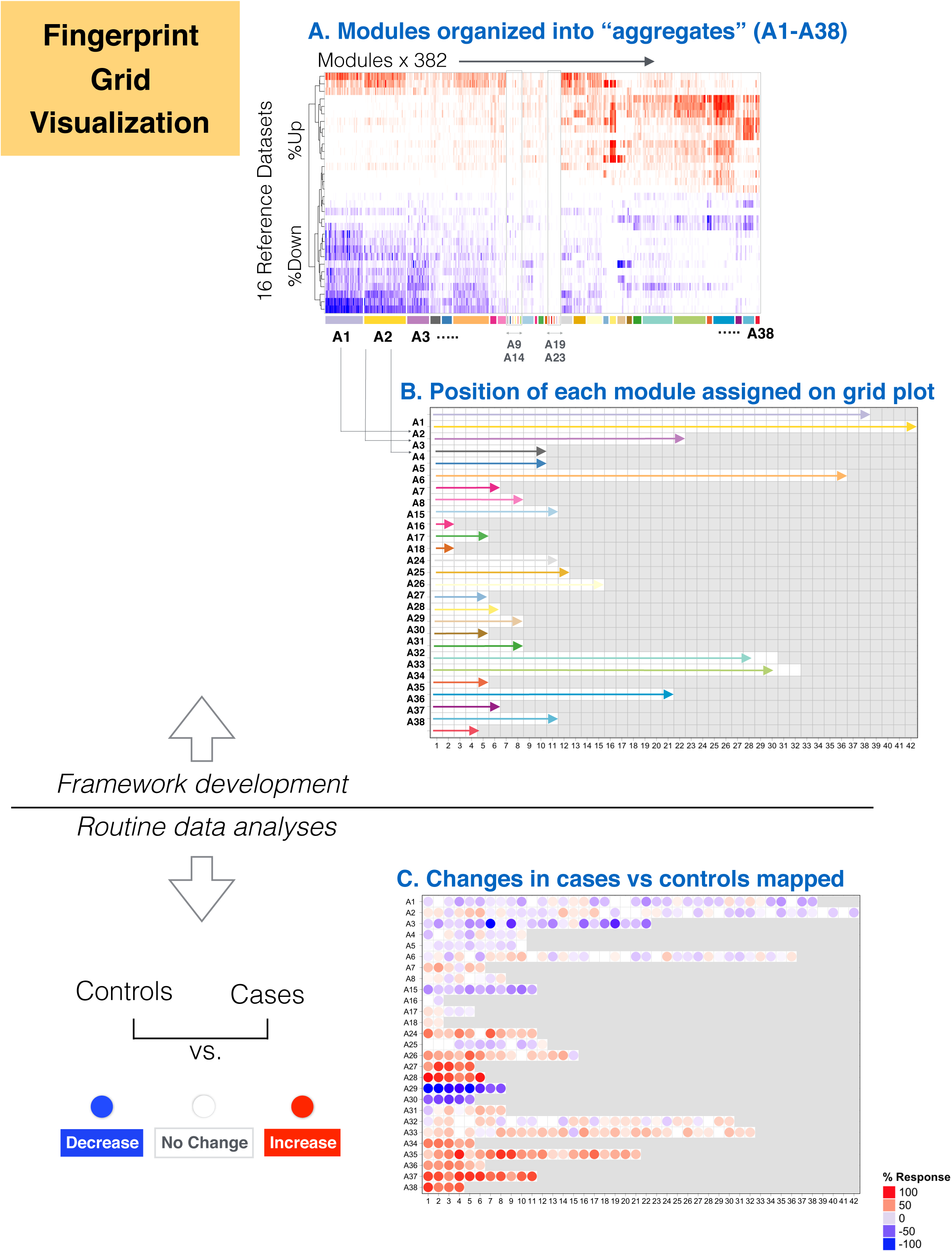
The development of module fingerprint grids. A. The modules were arranged onto the grid as follows: the master set of 382 modules was partitioned into 38 clusters (or aggregates) based on similarities among their module activity profiles across the sixteen reference datasets (A1-A38). B. A subset of 27 aggregates comprising 2 modules or more in turn occupied a line on the grid. The length of each line was adapted to accommodate the number of modules assigned to each cluster. The format of the grid was fixed for all analyses carried out using this modular framework. C. Changes in transcript abundance at the module level were mapped onto this grid and represented by color spots of varying intensity.

**Figure 3:**
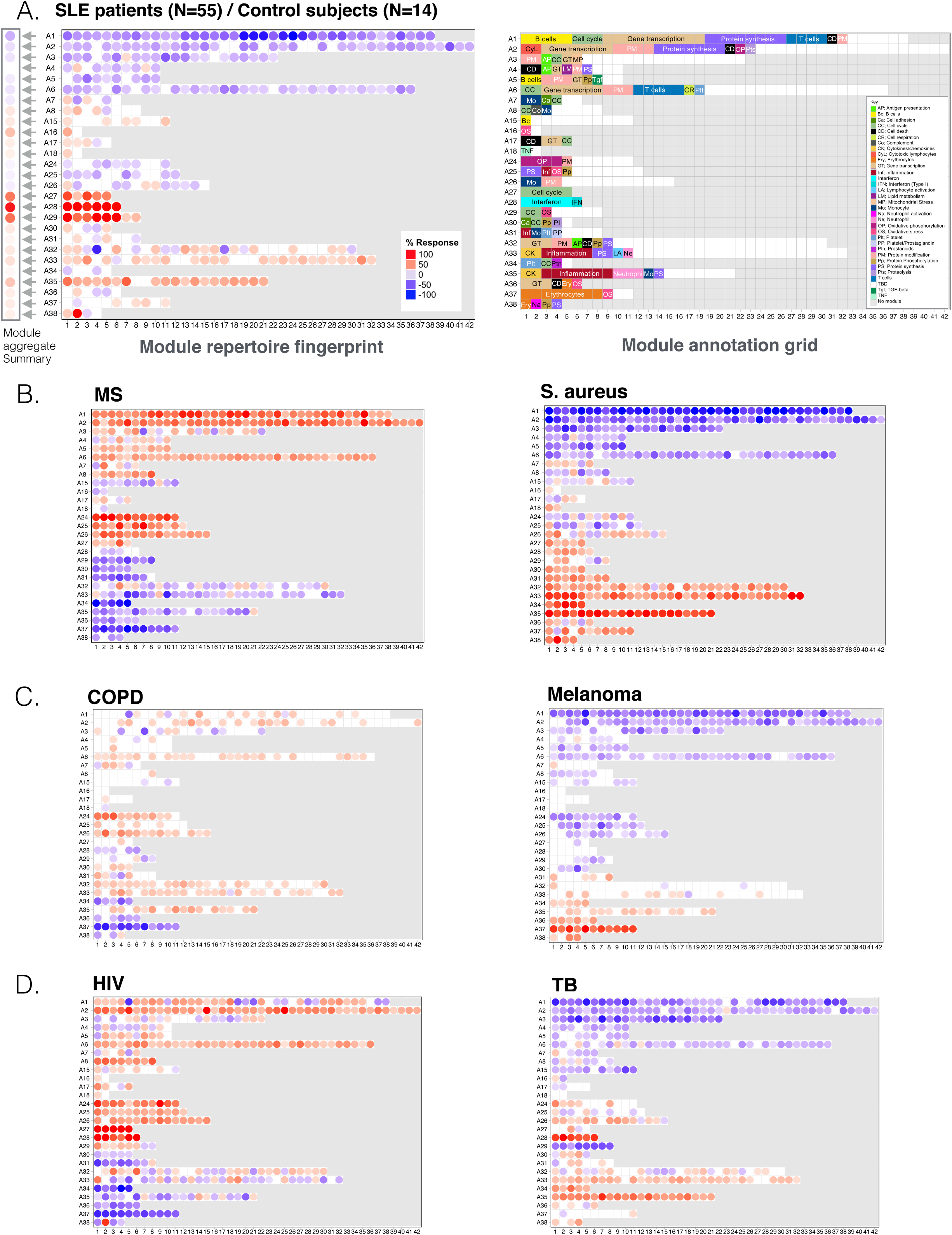
Fingerprint grid plots. A. Prototypical fingerprint grid plot: Changes in blood transcript abundance for patients with SLE compared to healthy controls are represented on a fingerprint grid plot for this illustrative use case. The modules occupy a fixed position on the fingerprint grid plots (see Figure 2). An increase in transcript abundance for a given module is represented by a red spot; a decrease in abundance is represented by a blue spot. Modules arranged on a given row belong to a module aggregate (here denoted as A1 to A38). Changes measured at the “aggregate-level” are represented by spots to the left of the grid next to the denomination for the corresponding aggregate. The colors and intensities of the spots are based on the average across each given row of modules. A module annotation grid is provided where a color key indicates the functional associations attributed to some of the modules on the grid (top right). Positions on the annotation grid occupied by modules for which no consensus annotation was attributed are colored white. Positions on the gird for which no modules have been assigned are colored grey. B-D. Fingerprint grid plots for additional reference datasets.

Overall, fingerprint grid plots can complement traditional heatmap representations. With the positions of modules on the grid being fixed it becomes for instance possible to rapidly identify the key changes associated with a given pathology or immunological state.

### Illustrative use case of fingerprint grid plot representations

We next illustrate the analysis and visualization approach described above with a use case. It will primarily focus on SLE, a disease which blood transcriptome signature has been well-characterized. Fingerprints of other reference disease cohorts employed for module construction will be included to provide additional context.

As mentioned, data interpretation is facilitated by tiered dimension reduction: the first vertical reading of the fingerprint grid permits visualization of changes across the aggregates, while the horizontal reading permits visualization of changes within an aggregate and across modules. As pointed out earlier, in the fingerprint grid, the number and intensity of the spots represents qualitative as well as quantitative differences. As an example, we compared the transcriptome profiles among 55 pediatric patients with SLE and 14 healthy control subjects (**Figure 3**). We identified an interferon-dominated signature (A28) accompanied by modules associated with cell cycle (A27 and A29, including antibody production). An increase in the abundance of modules associated with inflammation and neutrophils (A35) — a hallmark of the SLE transcriptome signature was also observed. These changes were accompanied by a decrease in transcript abundance, which was more apparent for some modules belonging to aggregates A1, A2 and A3 which are arrayed across the first three rows of the fingerprint grid. More specifically, for the module aggregate A1, the most marked decreases were observed for modules associated with protein synthesis (dark purple color at positions 1, 5, 11 and 19 on row A1).

It is possible to go a step further and “aggregate” the changes observed by row, thereby reducing the dimensions for a given dataset even further. In this case, we reduced the dataset from 382 modules to 27 “aggregates” (**Figure 3A**). Users can decide taking this extra step depends on the desired level of resolution: module-level or aggregate-level. For example, mapping changes at the aggregate level, the simplest framework possible, is optimal when looking at signatures in a broader context. But at the same time our earlier work showed that distinct interferon modules are biologically and clinically meaningful, also it can also be indicated to work instead at the module-level (11).

We generated fingerprint grid plots for each of the sixteen diseases or physiological states (**Supplementary file 3**); six module fingerprints are shown in **Figure 3** as an illustration. In brief, among these we found that blood transcriptome perturbations were most widespread in patients with MS and patients with a *Staphylococcus aureus* infection (**Figure 3B**) with opposing patterns of change. Changes associated with COPD or stage IV melanoma (**Figure 3C**) were most subtle but nonetheless distinct, with differences in transcript abundance compared to control subjects most visible for aggregates concerning oxidative phosphorylation, monocytes, inflammation (A24-A26), erythrocytes and neutrophil activation (A36-A38). Interferon signatures were a common trait (A28) in patients with SLE and patients with tuberculosis (TB) caused by *M. tuberculosis* infection (**Figure 3D**), while opposite patterns were observed for the cell cycle (A29). We also found differences in the intensities of sets of modules associated with inflammation between these two diseases (A33-A35).

Taken together, this use case demonstrates the use of “fingerprint” representations. Notably, fixing positions on the grid permits to overlay functional annotations associated with each modular signature, and the use reference collections of fingerprints for comparative interpretation. However, while this representation can be used as a complement, it altogether does not replace the more traditional heatmap representations.

### In depth functional annotation of fixed transcriptional module repertoires

Based on earlier iterations, the expectation is for this repertoire to be of use over a period spanning at least 5-6 years. We thus put great effort into functionally annotating the repertoire. We followed two main annotation approaches: (1) concurrently running ontologies, pathways or literature-term profiling analyses; and (2) determining for each module the patterns of gene expression for select reference transcriptome datasets. We compiled the resulting information for all 382 modules and have since made it available via an interactive web application Here, it is possible to zoom in and out, determine spatial relationships and interactively browse the very large compendium of analysis reports and heatmaps generated as part of our annotation efforts. (**Figure 4**; Links for each aggregate are listed in **Table 3**.; A demonstration video is available: https://youtu.be/8ajESET2mqI) Below, we describe briefly how we conducted our annotations

**Table 3:**
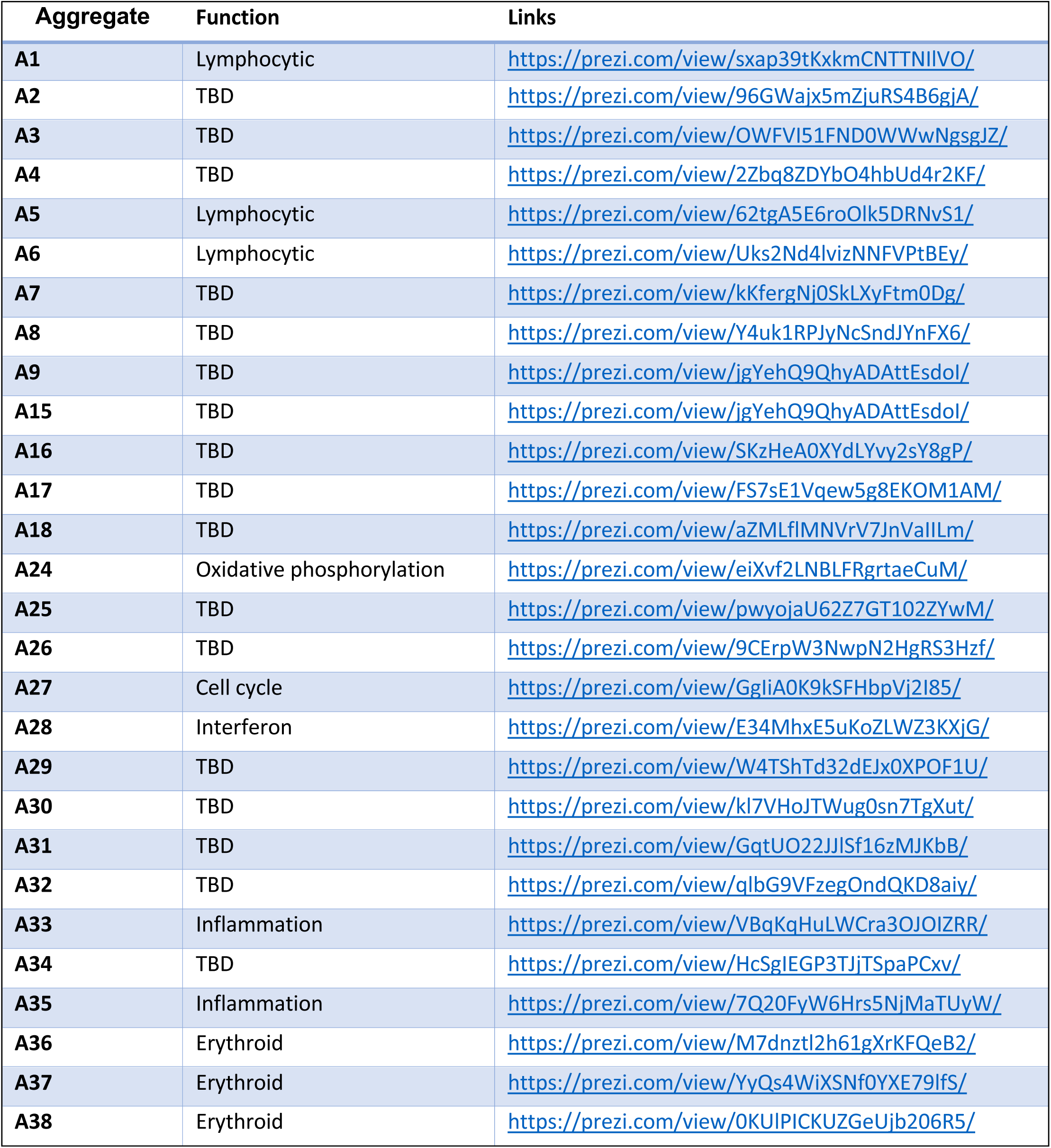
Links to module aggregates annotation pages.

**Figure 4:**
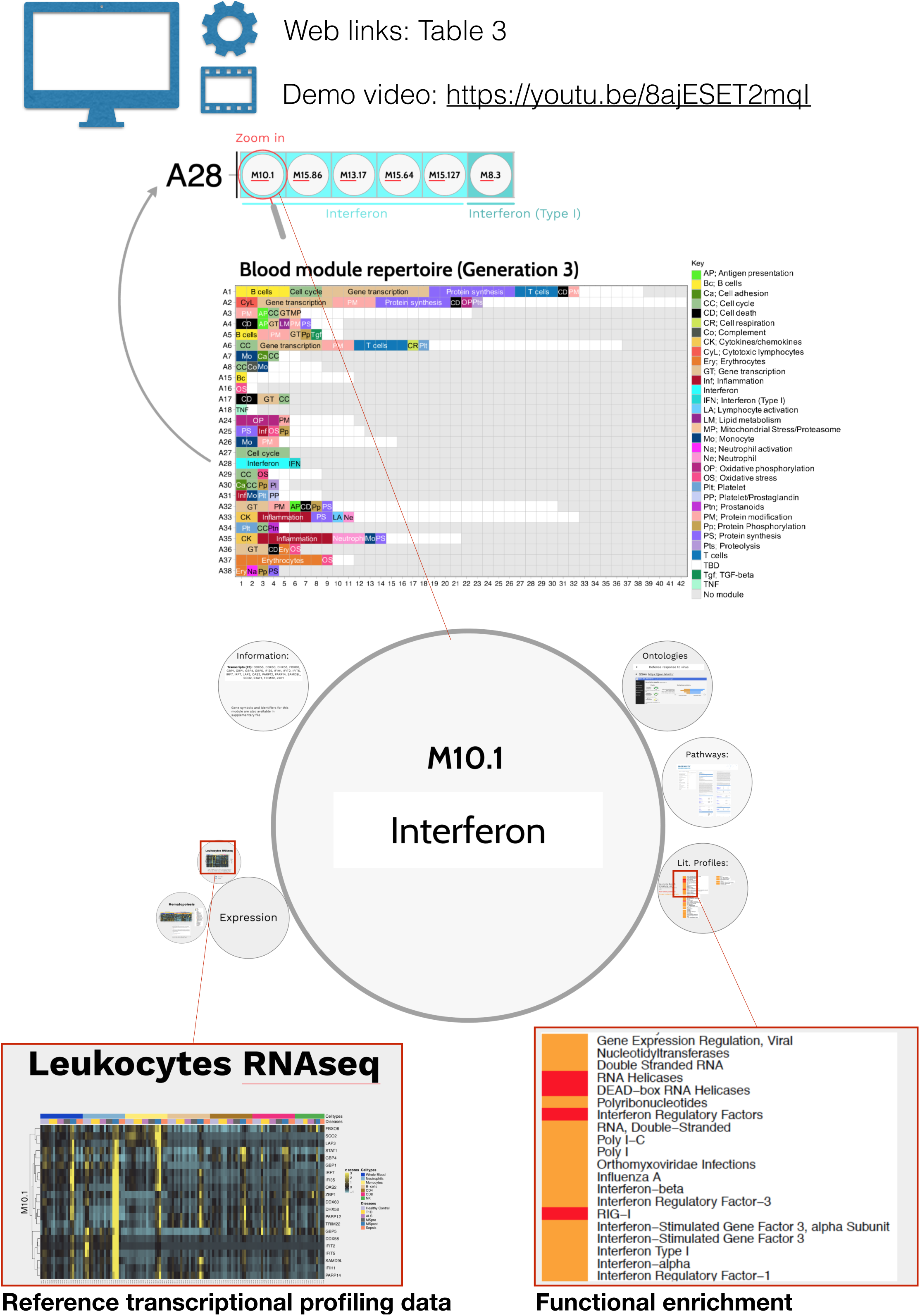
Functional annotation of the transcriptional module repertoire. An interactive application is available to explore the 382 modules comprising the blood transcriptome repertoire. A gene list, along with the ontology, pathway, literature term enrichment and transcriptional profiling data for reference transcriptome datasets (circulating leukocyte populations, hematopoiesis) is provided for each module. Zoom in and out functionalities for close-up examinations of the text and figures embedded in the presentation is possible. Web links providing access to modules within a given aggregate are listed in Table 3. For a demonstration video, please visit: https://youtu.be/8ajESET2mqI.

(see also Experimental Procedures).

*Step 1 Functional profiling:* We conducted gene ontology profiling for each of the 382 modules using DAVID (12), GOTERM, and GSAn (13). GSAn interactive reports were uploaded to a custom web portal (https://ayllonbe.github.io/modulesV3/index.html). We also performed pathway enrichment analyses using KEGG, Biocarta and Ingenuity Pathway, as well as literature term enrichment with Literature Lab. We synthesized and compiled the analyses (**Supplementary File 2**) and used them to identify convergences and attribute functional titles to the different modules. Functional titles could not be attributed in all cases, due to a lack of convergence or poor enrichment in one or several of the analyses.

*Step 2 Expression patterns in reference transcriptome datasets:* We used transcriptome datasets as a reference to improve characterization and biological interpretation of the module framework. Two different datasets were used. The first was contributed by Novershtern *et al*. and comprised the transcriptome profiles of 38 human hematopoietic cell populations (14). The second was contributed by Speake *et al*. and comprised the RNAseq profiles of six circulating leukocyte populations from patients with various immune-associated diseases (15). We generated heatmaps for each module to display the abundance patterns of the constitutive transcripts for each dataset.

Overall, this resource serves two purposes. First, it provides access to information that is necessary for interpreting transcriptome fingerprints generated via module-level analyses. Second, it helps us to improve the attribution of functional titles and roles to the different modules and aggregates. Indeed, although the transcriptional module repertoire is fixed over time, we anticipate that the functional annotations will continue to evolve over its lifespan.

**Measuring inter-individual variability for the molecular stratification of patient cohorts** The analysis and visualization steps presented so far focused on characterizing differences between groups of subjects (e.g. cases and controls). However, it is also important to characterize heterogeneity among groups of patients since inter-individual variability can serve as a basis for the definition of molecular phenotypes and patient stratification.

Within each module, and for each individual subject, we used fixed cutoffs to count the number of transcripts that increase, decrease or do not change in abundance compared to a baseline value (e.g. absolute fold change in expression and absolute difference in expression vs. average of control samples). The percentage of differentially expressed genes for each module are then computed. These percentages are equivalent to values derived from group comparisons, except that they are derived for each individual sample.

The sepsis cohort comprised in the reference dataset collection was used to illustrate how this approach can be employed for assessment of inter-individual variability for a given pathology (**Figure 5**). Changes in transcript abundance across was found to be highly consistent across patients for some module aggregates. This was for instance the case of aggregate A1 (broadly associated with lymphocytic cells/responses), with consistent decreases in transcript abundance observed across patients. Conversely, consistent increases were observed for modules comprising aggregate A35 (broadly associated with inflammatory neutrophil responses). In this case differences were observed in the intensity of the response. Other module aggregates, however, showed more mixed responses, which was the case of A37 (erythroid cells), A33 (functional association to be defined) or A28 (interferon response).

**Figure 5:**
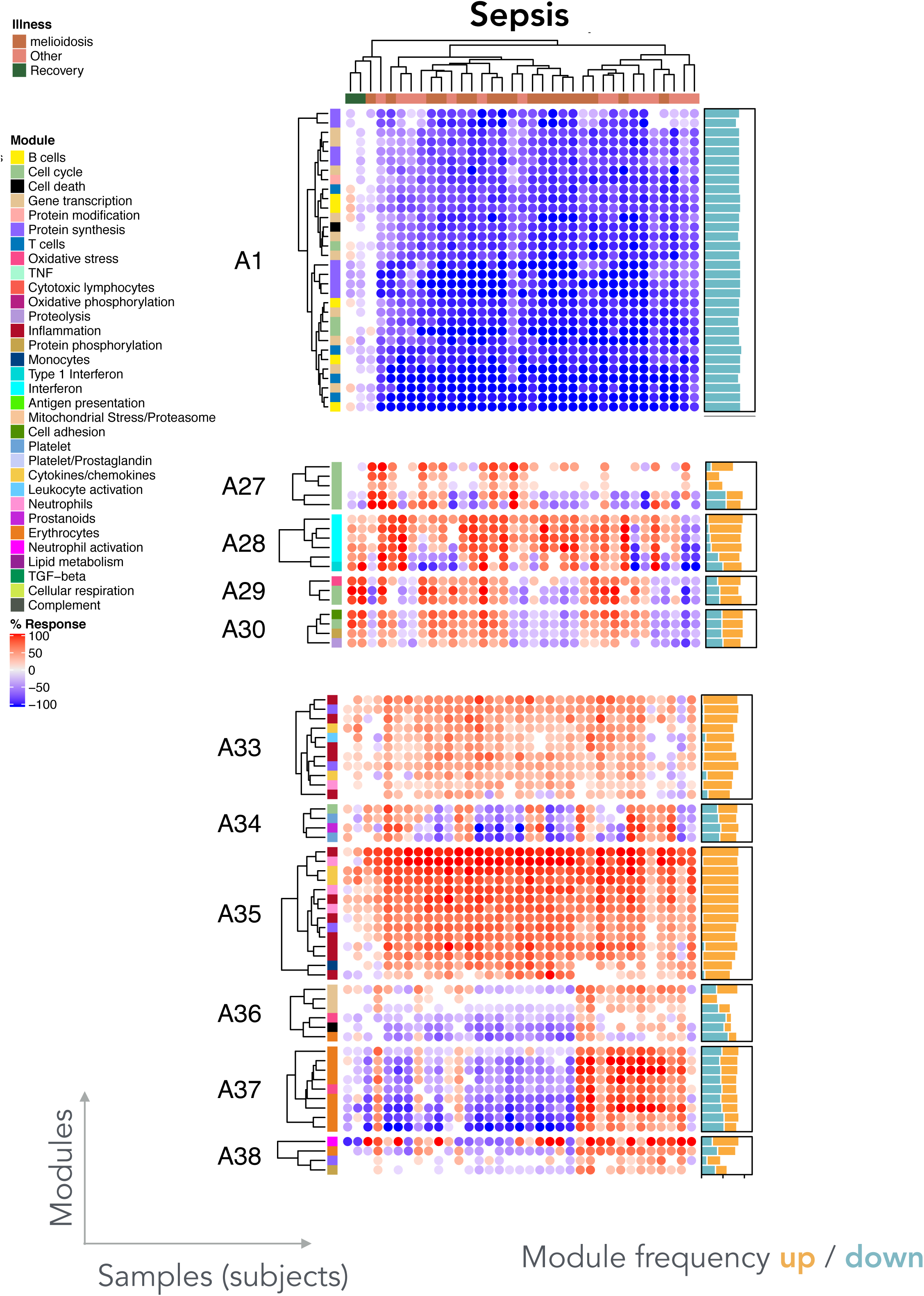
Individual-level module heatmap. Changes in transcript abundance were determined at the individual level across all modules constituting the repertoire. These changes are represented on a heatmap, where an increase in abundance of a given module is represented in red, and a decrease in abundance is represented in blue. The subjects are organized as columns and the modules as rows. The ordering on the heat map was determined by hierarchical clustering.

A web application was developed as a resource to explore the inter-individual differences for a given disease, module aggregates or a combination of aggregates (https://drinchai.shinyapps.io/dc_gen3_module_analysis/#; video: https://youtu.be/y7xKJo5e4). This application permits the generation of fingerprint grid plots and heatmaps representing module aggregates activity across the 16 reference datasets (**Figure 6**).

**Figure 6:**
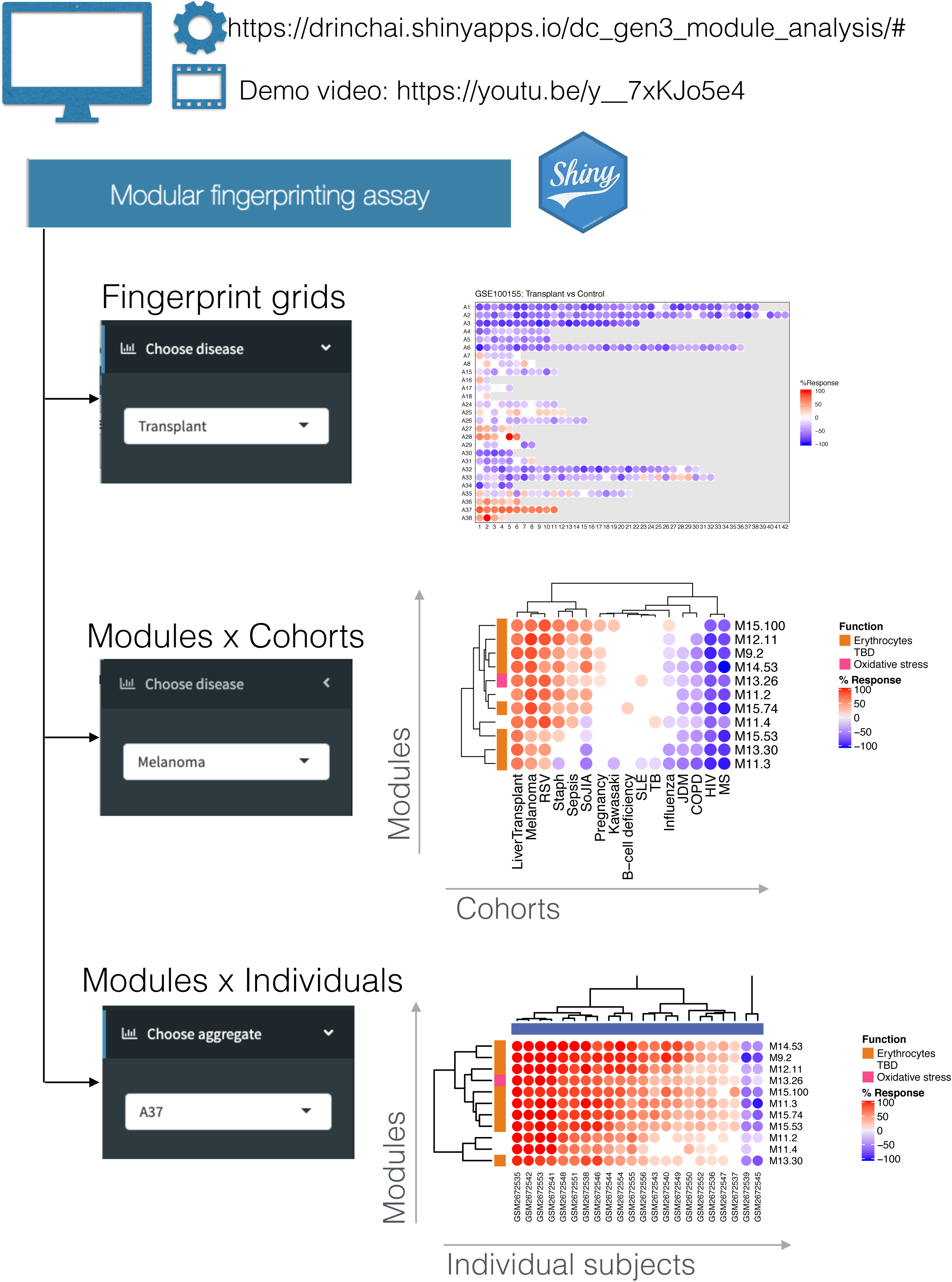
Web application to visualize multi-tiered module fingerprinting. An application was developed to explore the changes in transcript abundance at the module level across the 16 reference datasets used to construct the repertoire. Three types of plot can be displayed and exported: 1) fingerprint grids; 2) module heatmaps displaying changes in abundance in modules comprising a given aggregate across the 16 reference datasets; and 3) module heatmaps displaying changes in abundance in modules comprising a given aggregate across individuals constituting a given dataset. For accessing the application, please visit: https://drinchai.shinyapps.io/dc_gen3_module_analysis/#. For a demonstration video, please visit: https://youtu.be/y_7xKJo5e4.

Altogether, as illustrated by the sepsis use case provided here, deriving individual modular fingerprints can provide a means to achieve molecular stratification of patient cohorts. However, the biological and clinical relevance of such distinct molecular phenotypes would still remain to be determined in follow on analysis steps.

### Profiling the abundance of A28 interferon-inducible genes at the aggregate-level across reference patient cohorts

The earlier sections dealt first with the approach implemented for the construction and characterization of the fixed transcriptional module repertoire. Analysis and visualization strategies for both group-level and individual-level comparisons were presented next. Now we would like to go over a use case focusing on the dissection of changes in abundance for module aggregate A28 (interferon responses). We first start here from the highest possible perspective, examining changes in abundance for the A28 aggregate across reference disease cohorts.

A heatmap was derived showing patterns of abundance of a subset of 27 module aggregates comprising two or more modules across 16 health states (**Figure 7A**). The first order of separation grouped acute HIV infection, MS, juvenile dermatomyositis and COPD in one cluster. The remaining 14 states grouped into a second cluster, with RSV infection presenting as an outlier. The main trend driving the dichotomy between the first four diseases and the rest was an overall suppression of modules associated with inflammation and/or myeloid cell responses (A34-A38), accompanied by an increase in modules corresponding to aggregates associated, in part, with lymphocytic responses (A1-A8). The factors underlying these two distinct, “overarching” signatures are unclear; diseases belonging to either group can exhibit marked interferon signatures (e.g. acute HIV infection in one cluster, and SLE or influenza infection in the other cluster). The circle plots shown on this figure illustrate at a more granular module and gene-level the changes that are being represented by spots on this heatmap. The gene composition for each of the six module is shown in one of those circle plots (**Figure 7B**). The other represent which of the genes among those modules are changed when comparing patients with infection to their respective control groups (**Figure 7C**). Finally, we added for reference circle plots showing patterns of *in vivo* response of A28 genes to administration of IFNα to patients with hepatitis C infection or of IFNβ to patients with multiple sclerosis [transcriptome profiling data were made publicly available by the authors (16,17)] (**Figure 7D**). The plots on this figure show that changes observed at the aggregate level are not always distributed evenly across the six modules constituting aggregate A28, and in turn of genes constituting each of the modules. The response to type I interferons was dominated by a disproportionate increase in abundance of transcripts constituting M8.3 and M10.1. Transcripts forming M15.86, that showed very little changes in response to those treatments, were on the other hand markedly increased during acute HIV and influenza infection (**Figure 7C**). It is thus possible that this gene set be more specifically induced by interferon gamma. Interferon responses were weaker among RSV patients compared to these two other viral illnesses. Similarly, in the context of bacterial infection the interferon response was most marked in response to TB infection, which is consistent with the literature (8). A large proportion of adult patients comprising the sepsis cohorts were infected with *Burkholderia pseudomallei*, the intracellular bacteria responsible for melioidosis that also tends to induce higher interferon responses (8). Changes in abundance for transcripts constituting A28 in the context of autoimmune or inflammatory diseases is shown in **supplementary Figure 2**.

**Figure 7:**
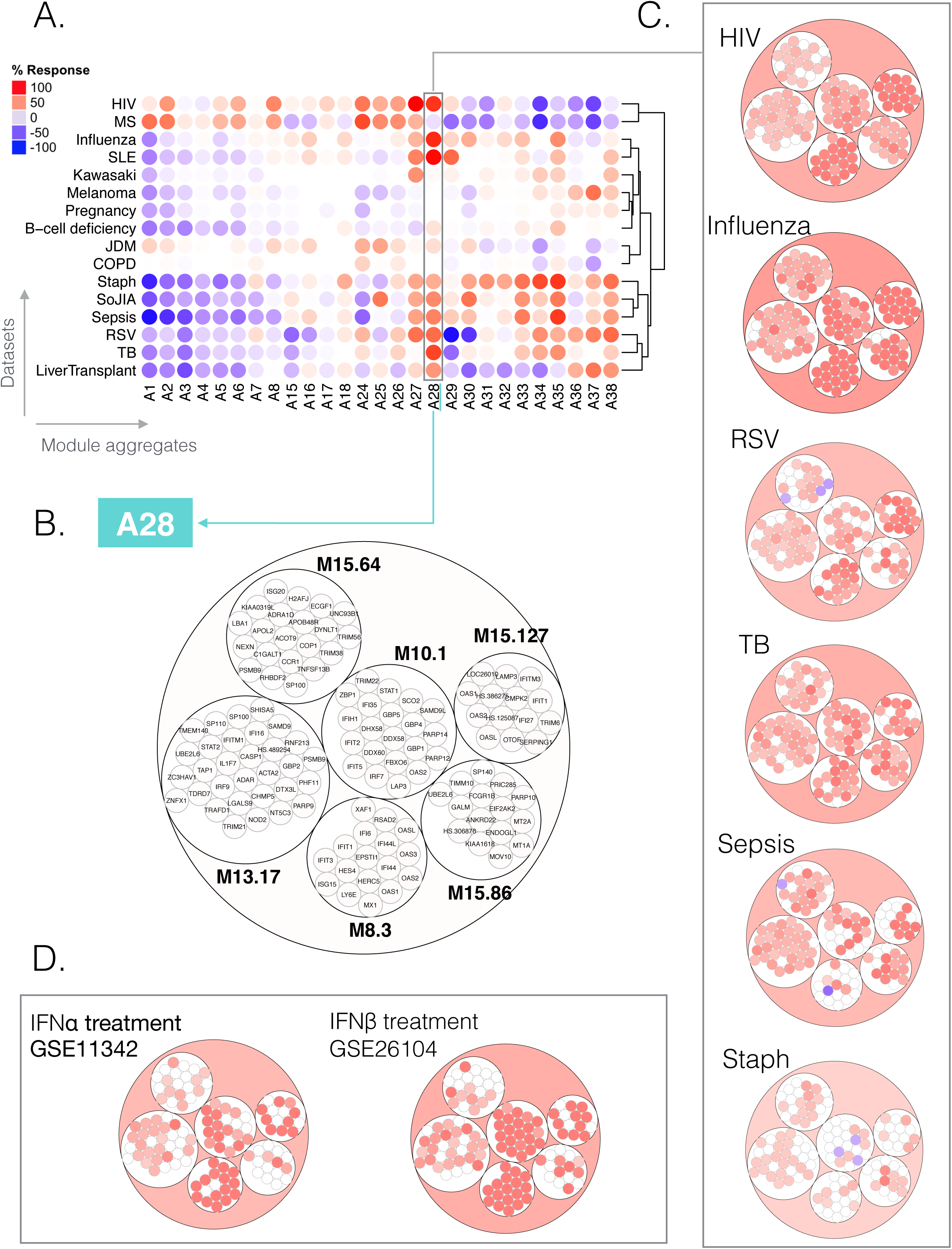
Module aggregate abundance patterns across the 16 disease or physiological states. **A. Patterns of changes in transcript abundance at the aggregate and cohort levels.** Each column on the heat map corresponds to a “module aggregate”, numbered A1 to A38. Modules A9- A14 and A19-A24 were excluded as they each only included one module. Each row on the heatmap corresponds to one of the 16 datasets used to construct the module framework. A red spot on the heatmap indicates an increase in abundance of transcripts comprising a given module cluster for a given disease or physiologic state. A blue spot indicates a decrease in abundance of transcripts. No color indicates no change. Disease or physiological states were arranged based on the level of similarity in the patterns of aggregate activity, determined via hierarchical clustering. **B. Representation of the modules and genes constituting aggregate A28**. The circle plot represents the six modules constituting aggregate 28, and the transcripts constituting each of the modules. Some genes on the Illumina BeadArrays can map to multiple probes, which explains the few instances where the same gene can be found in different modules. **C. Patterns of changes in transcript abundance at the module-level and gene-level for aggregate A28**. The circle plots illustrate the changes at the gene-level for this aggregate for 6/16 datasets. The position of the genes on each of these plots is the same as shown in panel B. Genes for which transcript abundance is changed are shown in red (increase) or in blue (decrease). **D. Patterns of changes in transcript abundance at the module and gene level for aggregate A28 in subjects treated with IFNα or IFNβ treatment** The circle plots show changes in abundance for A28 transcripts in patients with hepatitis C infection treated with IFNα [GSE11342 (16)] or patients with multiple sclerosis treated with IFNβ [GSE26104 (17)].

The visualization schemes employed here already affords some new perspectives about the contribution of different modules and genes to the overall aggregate-level interferon responses. However other variation in the selection of variables and granularity levels are possible and will be explored next.

### Profiling the abundance of A28 interferon-inducible genes at the module-level across reference patient cohorts

Another perspective was gained next by plotting changes of A28 genes across the same reference cohorts but at the more granular module-level, rather than at the aggregate level. It was also at this level that we chose to compare literature keyword enrichment profiles for each of the A28 module.

Functional enrichment analyses showed that all six modules in this aggregate were associated with the interferon response (see https://prezi.com/view/E34MhxE5uKoZLWZ3KXjG/). The heatmap (**Figure 8A**) showing literature enrichment profiles highlighted keywords associated with viral pathogens (“hepatitis”, “herpes” or “influenza”), as well as host-derived and pathogen-derived molecules (“RIG-I”, “Interferon”, “Interferon, “double stranded RNA”). From the six modules, four seemed to be “core” interferon modules (M15.86, M10.1, M8.3, M15.127), while the remaining two (M13.17, M15.64) were associated to the interferon literature to a lesser degree. These latter two modules were more strongly associated with the Herpes simplex virus than the other four modules while the four core modules were preferentially associated with hepatitis. The heatmaps available via the Prezi interface, which depict the gene expression profiles for the A28 modules provide additional perspectives. For example, the dataset from Speake *et al*. comprised samples of patients with MS collected immediately before and 24 h after the administration of their first dose of interferon-β (**supplementary Figure 3** [GEO ID GSE60424); (15)]. Despite the small number of subjects in this category, we observed a clear pattern of response to interferon *in vivo* across all six modules. This observation confirms the functional associations obtained from enrichment analyses.

**Figure 8:**
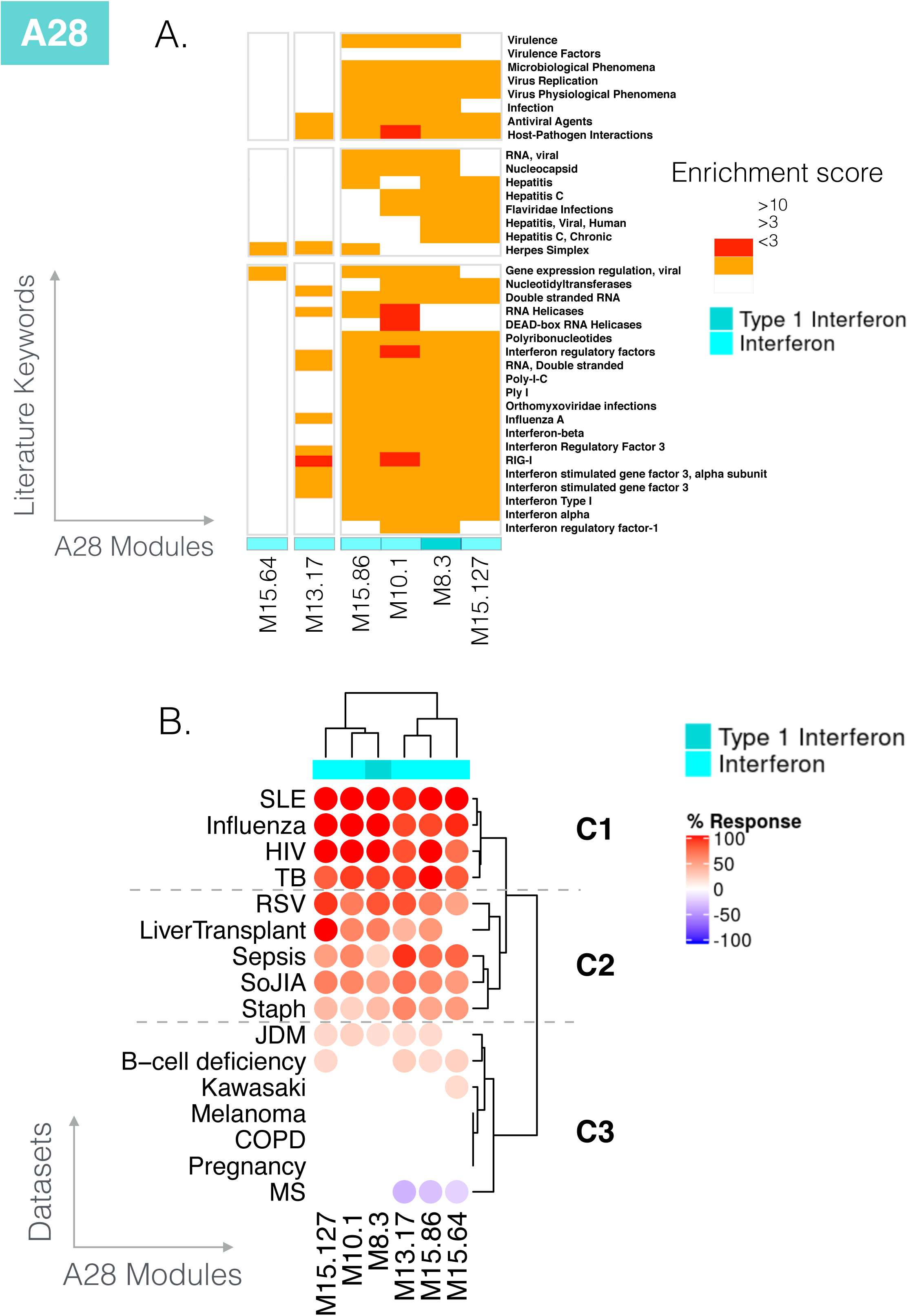
Literature profiles and patterns of changes in abundance across reference datasets for the modules comprising aggregate A28. A. Functional annotation by literature profiling: A portion of a heatmap comprising 382 modules organized as columns, and literature terms organized as rows. The six modules shown are associated with the consensus annotation “Interferon” or “Type 1 Interferon”. The clusters of keywords associated with those modules are consistent with this annotation and provide added granularity to the module repertoire for functional profiling and interpretation. **B. Changes in abundance across 16 reference datasets:** the heatmap (top) represents the changes in abundance of transcripts constituting the six modules comprising aggregate A28 (columns). The modules are functionally associated with interferon responses. The 16 reference datasets are arranged as rows corresponding to different health states. The columns and rows were arranged by hierarchical clustering. Such heatmaps can be accessed and exported for all 16 datasets and 38 module aggregates using the web application: https://drinchai.shinyapps.io/dc_gen3_module_analysis/# (under the “MODULES x STUDIES” tab).

Next, we examined the degree of changes for the six interferon modules across the 16 input datasets (**Figure 8B**). The first cluster showing the highest induction levels comprised SLE, influenza infection, HIV infection and active *M. tuberculosis* infection (noted as C1 on the figure). Interferon has antiviral properties; therefore, it is no surprise to see viral infections included in this set. Blood transcriptome profiling studies conducted nearly 20 years ago identified interferon responses in SLE pathogenesis (18,19). More recent profiling also revealed the prominence of this signature in patients with tuberculosis, which contrasts with findings made in other bacterial infections (6). The second cluster (noted as C2) comprised diseases with an “intermediate” level of interferon responses, including RSV infection, sepsis caused by *Staphylococcus aureus* in pediatric patients and a range of bacterial pathogens in adults, SoJIA and liver transplant recipients receiving maintenance immunosuppressive therapy. The illness conferred by RSV infection has a clinical presentation very similar to that of influenza in pediatric patients. However, studies suggest that interferon responses may be partially inhibited by RSV (20,21).

The final, third cluster was formed by pathologic and physiological states where an increase in abundance of interferon-inducible genes was either modest or inexistent. This cluster included patients with juvenile dermatomyositis and B-cell deficiency (low levels), Kawasaki disease, melanoma, COPD, pregnancy (no increase), and individuals with MS (apparent decrease). The latter observation presents some interest since one line of treatment for multiple sclerosis is consists in administering interferon beta. This should as a result compensate the defect observed here in treatment-naïve individuals.

Taken together, this closer examination of annotations for A28 modules and patterns of changes in abundance in different reference datasets provided a clearer picture of its biological significance.

### Profiling the abundance of A28 interferon-inducible genes at the module-level across individual subjects

As described earlier analysis workflows have been developed to determine changes in transcript abundance at the level of individual subjects. This approach, which offers an even more granular perspective, was applied next for molecular stratification of patient cohorts using the six A28 interferon modules.

We show, for instance, that in cohorts where no changes are detectable via comparisons at the overall group level, a minority of patients in fact present the signature. This scenario indeed applies for the cohort of melanoma patients with 3 out of 22 subjects showing some degree of increase in abundance for the six A28 modules, while the majority showed little changes or decrease in abundance. At least four of the patients showed a marked decrease in abundance (**Figure 9A**). The observation carries potential biological and clinical significance, as interferon activity in patients with melanoma has been previously associated with disease outcomes (22,23). Pathologies with an intermediate A28 signature might induce responses in a somewhat higher proportion of subjects. This is the case of JDM patient cohort that comprises a cluster of 11 patients our of 40 presenting with modular interferon signatures (**Figure 9A**). Of the 793 articles in PubMed mentioning “Juvenile Dermatomyositis” in their titles, 8 also mentioned “interferon”, which indicates that a role for interferon in this disease while not widely acknowledged has nonetheless been described (24). In diseases where the role for interferon is well described, such as influenza infection or SLE, increase in A28 modules is widespread. But it is nevertheless possible to find at least a few subjects in each cohort for whom an interferon signature is not present (**Figure 9A**). Notably in the case of SLE the proportion of interferon negative subjects tends to be higher in adult patient cohorts in comparison to pediatric patient cohorts as the one that is being used for illustrative purposes here. Stratification of an adult SLE cohort based on patterns of abundance of A28 modules is presented next.

**Figure 9:**
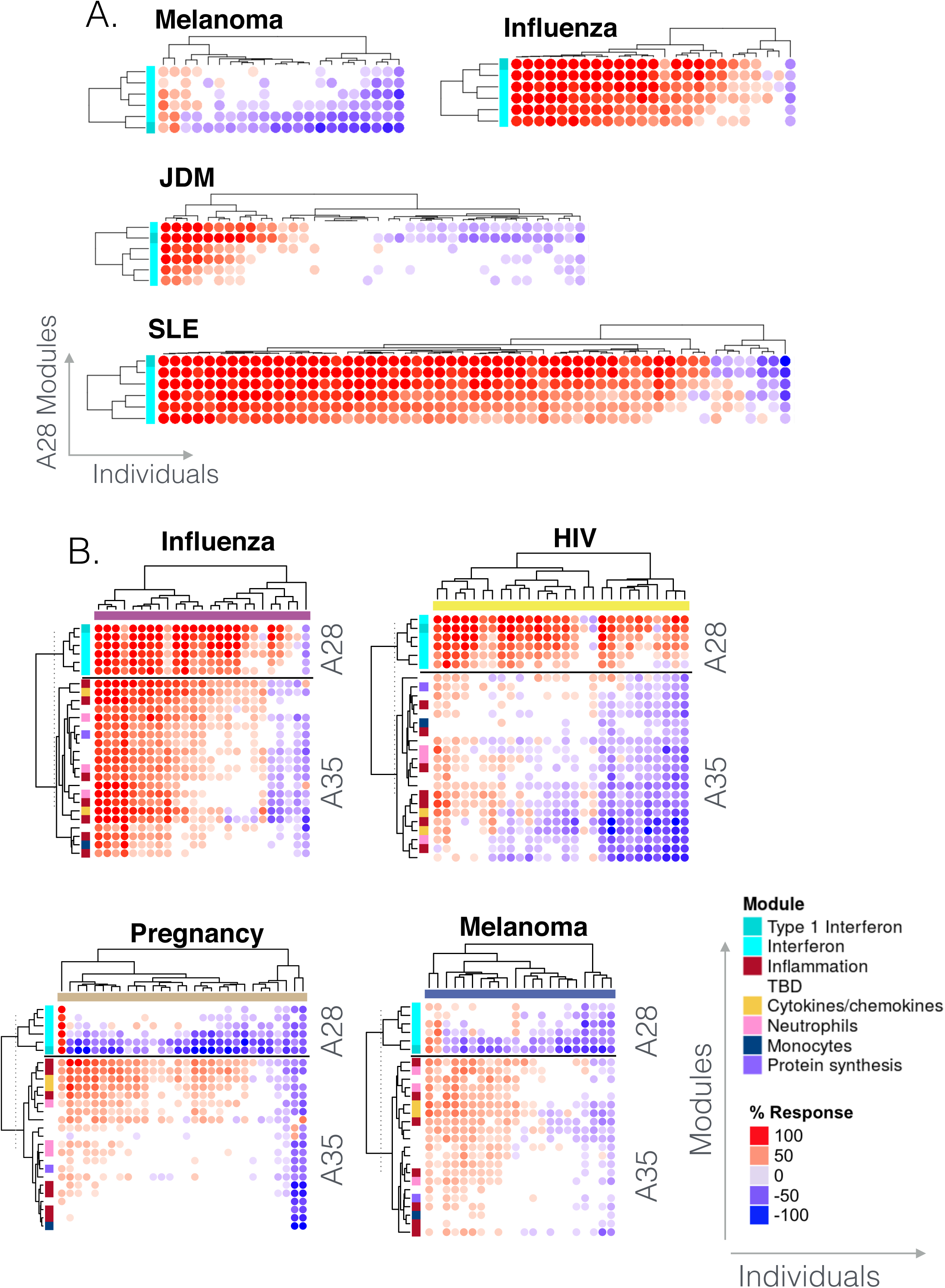
Abundance patterns across individuals. **A. Changes in abundance for A28 modules.** The heatmaps display the changes in abundance for the same six modules (rows) across individuals (columns) in four reference cohorts. The rows and columns on the heatmap are arranged based on similarities in abundance patterns. **B. Changes in abundance for A28 and A35 modules**. The heatmaps display the changes in abundance of six modules constituting aggregate A28 and 21 modules constituting aggregate A35 (rows) across individuals (columns) in four reference datasets. Functional annotations associated with different modules is indicated by a color code and corresponding legend. Such heatmaps can be accessed and exported for all 16 datasets and 38 module aggregates using the web application: https://drinchai.shinyapps.io/dc_gen3_module_analysis/# (under the “MODULES x INDIVIDUALS” tab).

We previously showed using our second generation repertoire that the interferon signature characterizing SLE comprises distinct “sub-signatures” at the module level (11). In this earlier work we observed a sequential increase for a set of three interferon modules (including M1.2, M3.4 & M5.12). M1.2 showed a higher degree of sensitivity, followed by M3.4 and then M5.12. Based on this finding, we could stratify SLE patients based on whether one, two or all three of those interferon modules were activated. By combining functional profiling with a reference dataset, we concluded that the modules responded differently to each interferon type: interferon-α induced an increase in the abundance of genes belonging to M1.2; interferon-β induced an increase in the abundance of genes belonging to M1.2 and M3.4; and interferon-γ induced an increase in the abundance of genes belonging to M5.12. Next, we sought to determine the equivalence between these three, second generation interferon modules and the six new, third generation interferon modules regrouped in aggregate A28. Based on gene composition, M8.3 and M15.127 mapped to M1.2 (inducible by both interferon-α and β), M10.1 and M15.86 mapped to M3.4 (inducible by interferon-β), and M13.17 and M15.64 mapped to M5.12 (inducible by interferon-γ). Notably, the latter two modules did indeed segregate from the other four on literature the enrichment profiling heatmap presented earlier (**Figure 8A**). We also used the six interferon modules to re-classify the adult SLE dataset profiled in our earlier study [GEO ID GSE49454 (11)]. The resulting clustering and stratification mirrored our earlier findings made using the three interferon modules (**supplementary Figure 4**). Overall these observations confirm that interferon “sub-signatures” may be employed for patient stratification. This may be relevant in the tailoring of biologics targeting interferon which are under development for the treatment of SLE. However, whether utilizing six modules from the new repertoire would necessarily be an improvement over the three from the second-generation repertoire that were employed previously remains to be determined.

Finally, we wanted to illustrate how further insights could be obtained by examining changes in A28 modules abundance at the individual level concurrently with that of other module aggregates. As an example, we present the changes for both A28 (interferon) and A35 (inflammation) modules (**Figure 8B**). By combining the two modular signatures, we could assess their relative contributions in a given cohort and identify distinct phenotypes: i.e. interferon or inflammation “positive”, “double positive” or “double negative”. We found the most contrasting patterns of change in transcript abundance in the sepsis and melanoma cohorts. Here, the abundance of interferon (A28) and inflammation (A35) modules almost uniformly increased in patients with sepsis. By contrast, we only observed increases in these modules in a minority of melanoma patients, with increases of inflammation modules being more widespread than that of interferon modules (∼50% compared to ∼10% of subjects, respectively). This finding was paralleled in pregnant women, who represent another immunosuppressive state, although increases in abundance of a subset of A35 modules appeared to be more prevalent in this group than in melanoma patients. Influenza infection was more similar to sepsis but with higher levels of interferon induction and a lower level of inflammation.

Taken together, the use case presented here illustrates the preliminary stepwise dissection of a given module aggregate and investigation of the underlying biological relevance. Such a process would be to be repeated for other module aggregates. While this is beyond the scope of the present work it is a process that the scalable annotation infrastructure that has been developed here will support. We indeed expect the range of reference datasets and functional profiling approaches available for interpretation to continue to expand. This work should also help determine to what extent subdivision of signatures in distinct modules is warranted.

## DISCUSSION

We have developed a new resource for the analysis and interpretation of blood transcriptome data. At its core is a fixed, transcriptional module framework. This framework has been constructed via co-expression analyses carried out across a wide range of diseases and physiological states. It constitutes the third iteration released by our group since 2008. Another blood transcriptome repertoire was developed and made available by our collaborators from Emory University [Li et al.: (9)]. While methodological differences exist for the construction of such frameworks the principle remains the same: i.e. utilization of reference datasets for constitution of reusable sets of transcriptional modules.

Collections of gene sets are commonly used for the interpretation of transcriptome profiling data [Gene Set Enrichment Analyses (GSEA): (25) (26)]. Such reference collections are very large, typically numbering tens of thousands of signatures. While they prove useful as a referential for functional interpretation, the majority of the sets are not derived via co-expression analysis. They would therefore not be sufficiently coherent to serve as a basis for dimension reduction, nor were they meant to be used in this fashion. Conversely, the construction of the set of transcriptional modules that is presented here is entirely driven by co-expression, and is suitable for reducing dimensions, but is intended for a much narrower use (i.e. the analysis and interpretation of blood transcriptome data). Zhou and Altman recently described the development of another fixed transcriptional module repertoire via co-expression analysis and discuss the advantages of such approaches (27). In brief, they assembled a collection of 2753 public datasets that were deposited in the NCBI GEO that encompassed 97,049 unique transcriptome profiles. They used independent component analysis for resolving co-expression relationships which led to the identification of a set of 139 transcriptional modules. They went on to demonstrate the biological relevance of these sets and advantages their use presents for improving robustness of analyses, especially for smaller datasets. Altogether this article provides additional arguments in favor of the reuse of fixed transcriptional module repertoires, an approach that to date has not seen widespread use so far. The authors chose a very wide breadth of datasets as input and the resulting framework may prove rather generic since transcriptional regulation is largely cell and tissue-specific.

A key characteristic of fixed transcriptional module frameworks is that they are intended for reuse, often over long periods of time. One of the major implications is that greater amounts of effort can be dedicated to the annotation and development of *ad hoc* visualization tools. This is obviously not possible when new module repertoires are constituted each time a new dataset needs to be analyzed. And indeed, considerable annotation and development efforts have been undertaken to support this latest generation of blood transcriptional modules. This permits us in turn to make available an ecosystem of custom tools and resources that were not available for transcriptional frameworks which have been released before. This includes a compendium of functional profiling reports and reference transcriptional profiles that can be accessed via an interactive web application. In addition, we are releasing another web application for data browsing and multi-tiered visualization. And it should also be noted that this ecosystem is scalable and will continue to grow over time as more analyses are being conducted, custom scripts developed, and reference dataset profiles added to the annotation scaffold. It is illustrated by ongoing efforts that consist in compiling relevant aggregate-centric information for A28. This is with the intent of improving over time understanding of the clinical relevance and biological significance of this transcriptional signature (accessible by clicking the information icon on the A28 interactive presentation: https://prezi.com/view/E34MhxE5uKoZLWZ3KXjG/ and video demonstration: https://youtu.be/0j-kcE1tlXA). A bioinformatics analysis toolkit that comprises a collection of custom R scripts will also follow the publication of this report. Notably, earlier attempts were made by our group to develop such a support infrastructure in the past, albeit on a smaller scale. But maintaining such resources over long periods of time proved a challenge and these by now have gone offline through a combination of hardware failures, hacking or discontinued institutional support. Such issues should be addressed here in part through adoption of zero cost infrastructure (e.g. hosting of Shiny apps, Prezi presentations or YouTube videos is free as long as public access is maintained).

Finally, the resource that we are making available here has some limitations which are also worth noting. First, our framework is specifically designed to analyze and interpret human blood transcriptome profiling data. Analyses of other tissues or in the context of other species would require the development and use of separate frameworks. Indeed, we also developed fixed modular frameworks for various mouse tissues (28) and *in vitro* culture systems, including human whole blood (29) and human dendritic cells (30). Second, we generated our transcriptome datasets for module construction using Illumina BeadArrays, a technology that pre-dates RNA sequencing. Our choice to use this technology was based on data availability at our institute, and the decision to use data that was generated at a single facility over a relatively short period of time. An alternative approach would be to compile available blood RNAseq data that is publicly available and that also comprises appropriately matched controls. The coverage afforded by RNAseq data would likely result in somewhat larger modular gene sets. However, we consider it unlikely that a fundamentally different repertoire would be produced using RNAseq data, as this would assume that entire co-expressed gene sets were missed by microarrays. Third, our modular repertoire is not meant to constitute a ground truth. Indeed, some aggregates are clearly not homogeneous in terms of functionality; and heterogeneity also exists within modules. The attribution of functional annotation titles to modules also confers some degree of subjectivity. However, the repertoire does structure the data so that insights about the biological significance of such a modular signature can be obtained. As a result, while the framework being presented will remain fixed for at least the coming few years, it is likely that the functional annotation map will continue to evolve for the foreseeable future.

Going forward, future developments include, firstly, the design and implementation of targeted transcriptional profiling assays based on the current repertoire. Indeed, such repertoires constitute a useful framework for module-based target selection and the development of “Transcriptome Fingerprinting Assays” [a preprint is available: (31)]. Such assays are cost-effective, robust and scalable. Secondly, fixed transcriptional module repertoires may be leveraged for conducting meta-analysis. It was for instance employed as a basis to perform such an integrative analysis encompassing six independent RSV datasets encompassing 490 unique profiles [a preprint is available: (32)]. More recently, this resource has also been used in a different context to identify signatures associated with CTLA4 and combined CTLA4-PD1 blockade in peripheral blood of ovarian cancer patients [P249: (33)]. It may also be possible to extend the overall approach described here to develop similar interpretive transcriptomic frameworks for other biological systems.

## ACKNOWLEDGEMENTS

The authors would like to thank Quynh-Anh Nguyen, Kimberly O’Brien, Dimitry Popov, Michael Mason, and Cate Speake for technical assistance. We would also like to acknowledge Insight Editing London for assistance in editing the manuscript prior to submission.

This project has been funded in part with Federal funds from the National Institutes of Health under contract number U01AI082110. Authors affiliated with Sidra Medicine, a member of the Qatar foundation for Education, Science and Community Development, are fully supported by institutional funding. RJW is supported by the Wellcome Trust (203135, 104803), NIH (U01AI115940) and the Francis Crick Institute, which receives funding from the Wellcome Trust (FC10218), CR UK (FC10218) and UKRI (FC10218).

## AUTHOR’S CONTRIBUTIONS

Conceptualization: MCA, DR, NB, DC. Data curation: NB, MA. Visualization: DR, MT, DC. Analysis and interpretation: MCA, DR, NB, AAB, EW, MG, BAK, MT, DC. Resources: MT, MA, SP, PK, LC, NJC, AAB, FM, PT, TP, GK, ML, HR, AOG, MB, CB, ML, RJW, CG, GL, MC, JS, RT, FK, AM, OR, KP, VP, JB. Writing of the first draft: DC. Funding acquisition: GK, KP, VP, OR, JB, DC. Methodology development: MCA, DR, NB, EW. Writing review & editing: MCA, DR, NB, MT, EW, MG, BAK, MA, SP, PK AAB, FM, PT, LC, NJC, JTP, GK, AOG, MB, CB, RJW, CMG, ML, GL, DB, RT, FK, AM, OR, KP, VP, JB, DC. The contributor’s roles listed above follow the Contributor Roles Taxonomy (CRediT) managed by The Consortia Advancing Standards in Research Administration Information (CASRAI).

## FIGURE LEGENDS

**Figure S1:**
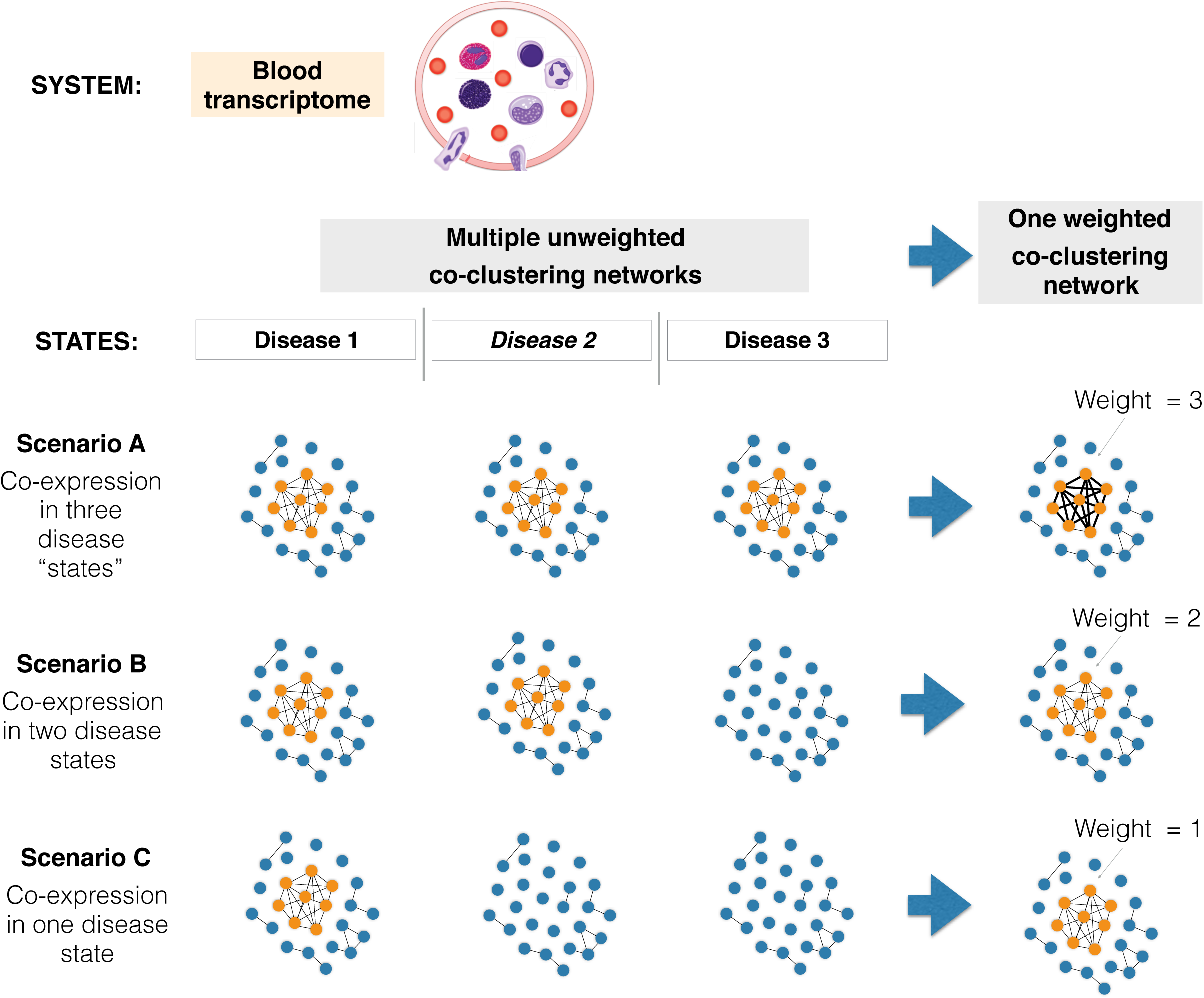
Construction of weighted co-clustering networks. Weighted co-clustering networks were used to construct the modular repertoires. These networks factor in differences in co-expression across different “states” of the biological system. For the blood transcriptome, these states would be different diseases or physiological phenotypes. Under Scenario A, the genes are co-expressed in all three disease states, so the weight attributed to the edges of the network is three. Under Scenarios B and C, co-clustering only occurs in two or one of the disease states, resulting in weights of 2 and 1, respectively.

**Figure S2:**
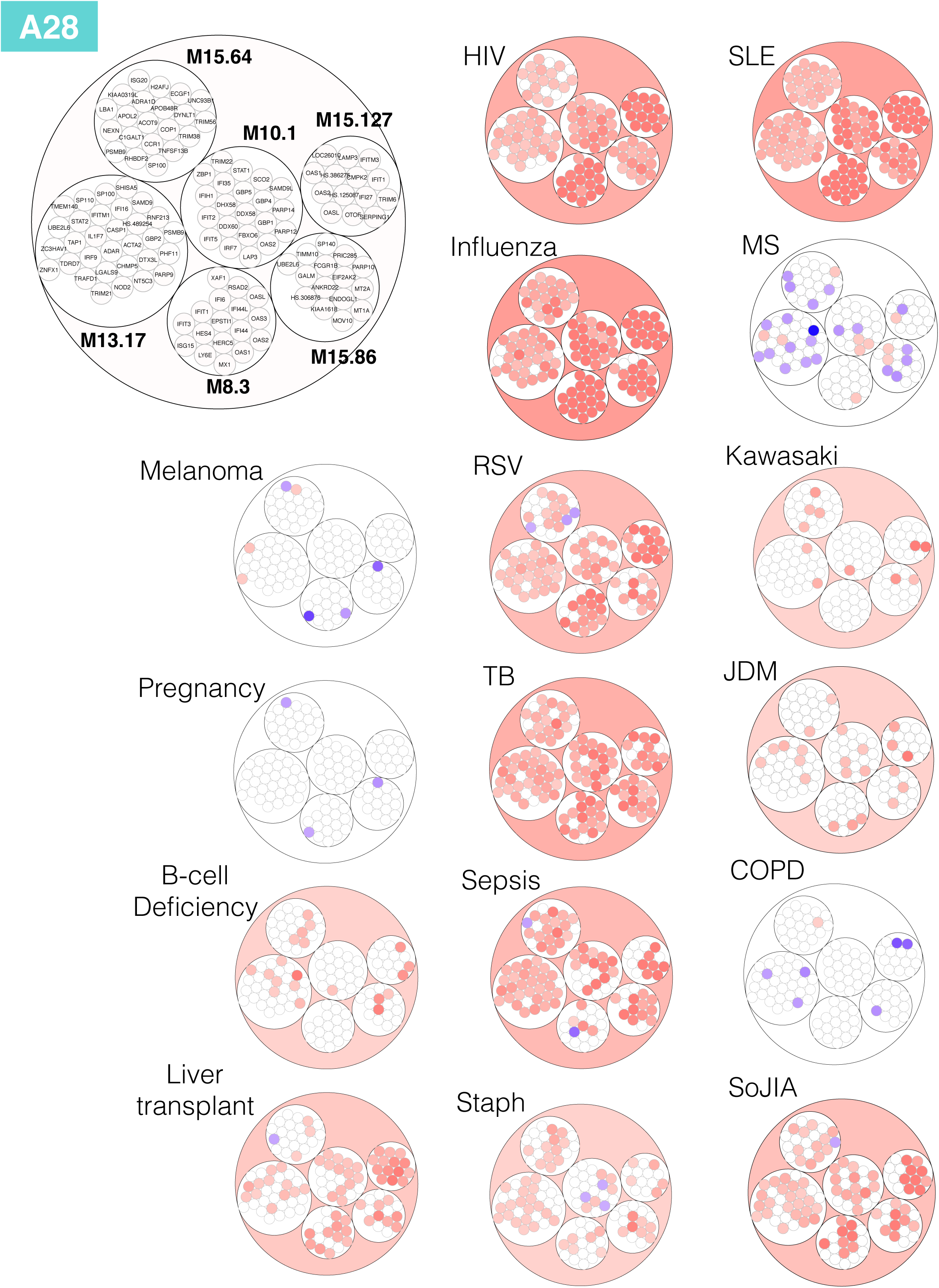
Patterns of changes in transcript abundance at the module and gene level for aggregate A28. The large circle plot represents the six modules constituting aggregate 28, and the transcripts constituting each of the modules. Some genes on the Illumina BeadArrays can map to multiple probes, which explains the few instances where the same gene can be found in different modules. Patterns of changes in transcript abundance at the module-level and gene-level for aggregate A28 are shown for all 16 reference datasets. The position of the genes on each of these plots is fixed. Genes for which transcript abundance is changed are shown in red (increase) or in blue (decrease).

**Figure S3:**
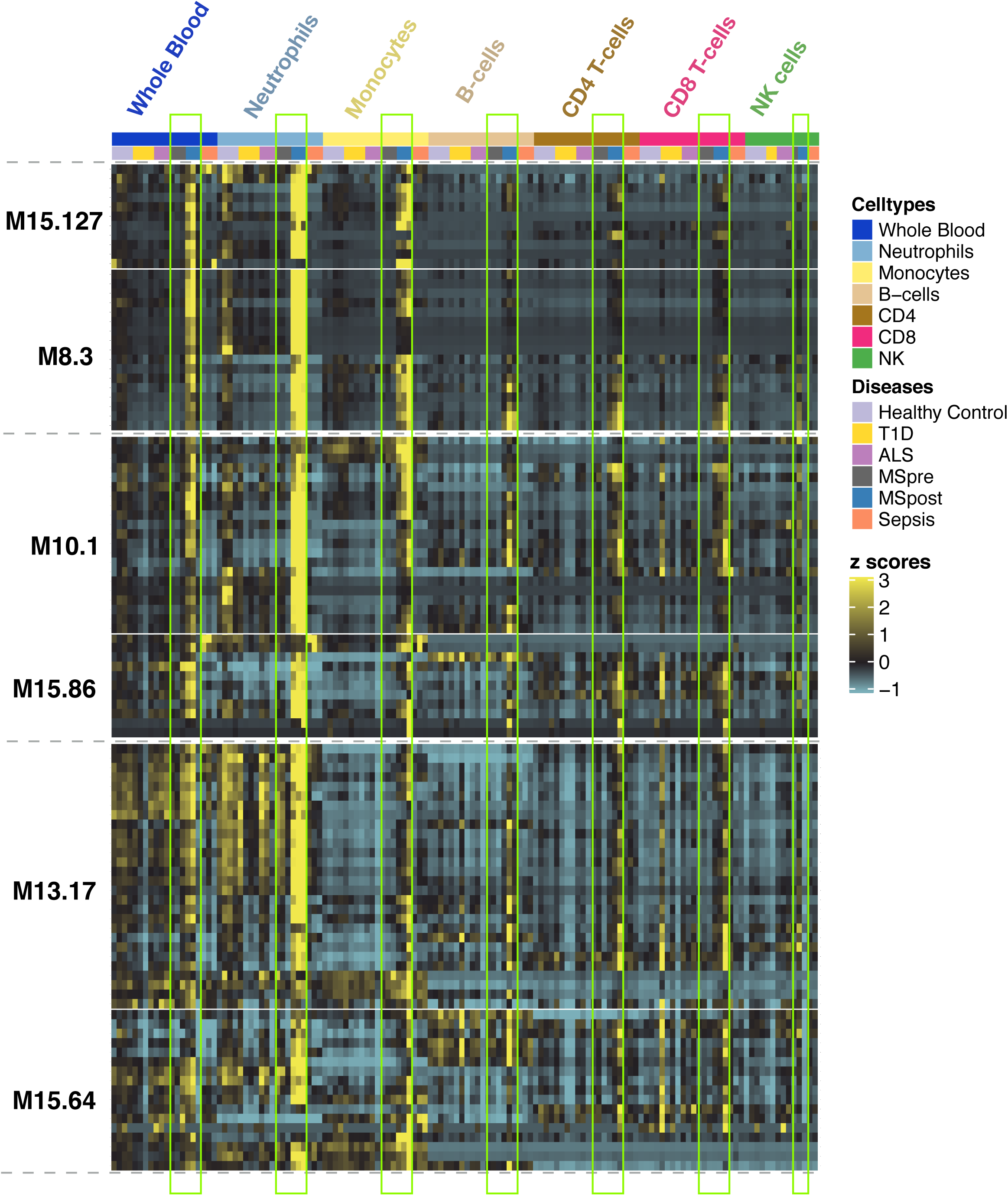
Expression patterns of genes constituting the six A28 modules (rows) across whole blood and purified circulating leukocyte populations (columns). This dataset was contributed by our group [Speake *et al*. (GSE60424)], and compared the RNAseq transcriptome signatures of six immune-cell subsets and whole blood from patients with various immune-associated diseases (color coding is provided to indicate cell populations and health status). The columns highlighted by the green boxes show the expression levels in patients with MS before and 24 h after the first dose of IFN-β.

**Figure S4:**
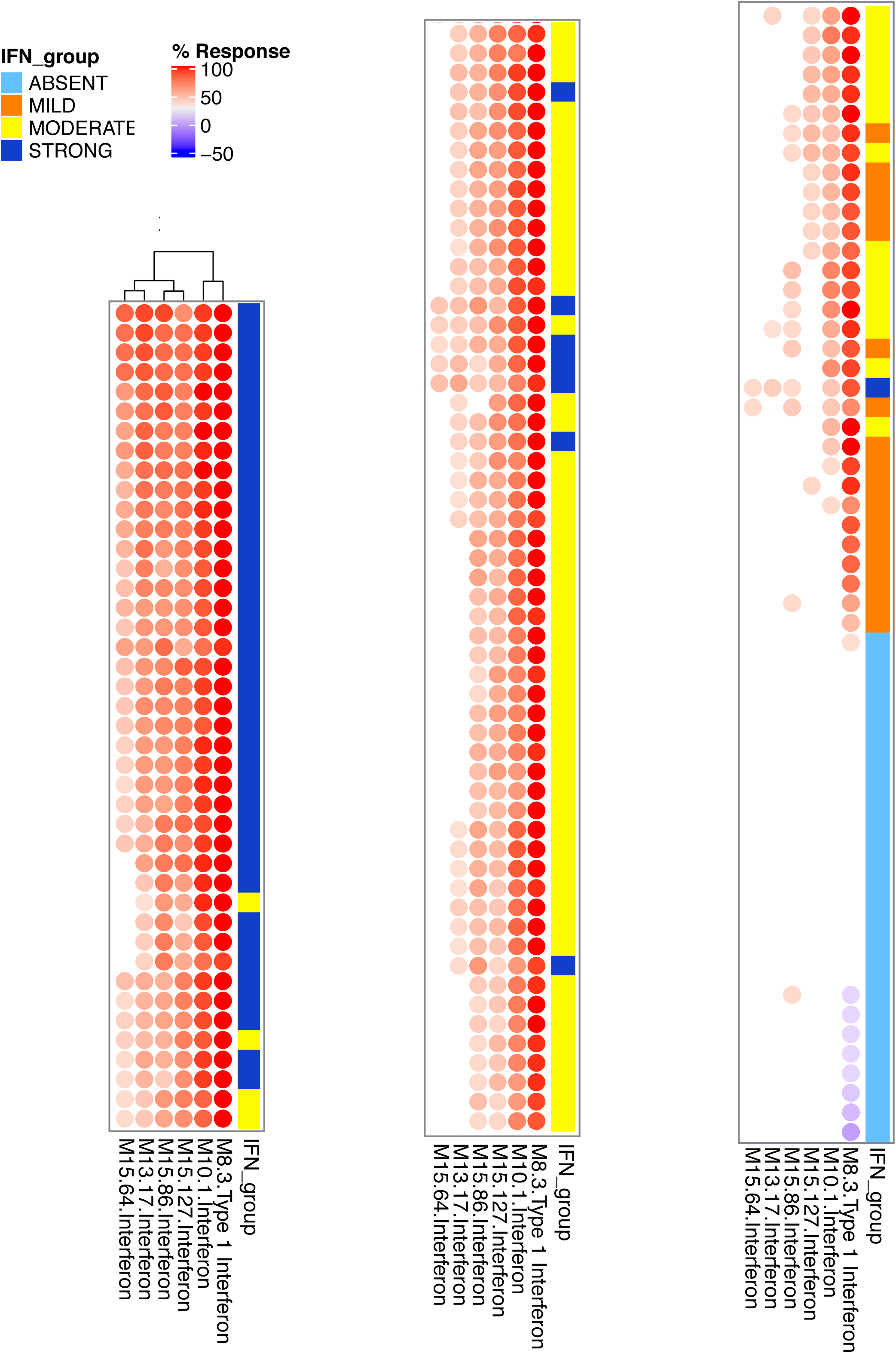
Abundance patterns of six interferon modules (A28) across individuals from an adult SLE cohort. The abundance levels for six interferon modules belonging to aggregate A28 (columns) is shown across a cohort of adult subjects diagnosed with SLE (rows). These subjects were part of the LUPUCE study: the dataset was contributed by our group [Chiche *et al*. (GEO ID GSE49454)] (11). Changes in transcript abundance were measured in reference to a cohort of healthy subjects included in the study. The colored label with interferon grouping information indicates the classification obtained by Chiche *et al*. based on three “second generation” modules.

## METHODS

### Study subjects

Gene expression datasets from 985 de-identified subjects from distinct cohorts were used for this work. Written informed consent was obtained from all participants. Studies were approved by Institutional Review Boards of the Baylor College of Medicine (COPD dataset: H-18029), the University of Texas Southwestern Medical Center and Baylor Health Care System (Influenza, RSV, S. aureus and Kawasaki disease datasets: UTSW #0802-447 BIIR #002-141), Saint Jude’s Research Hospital (B-cell deficiency), the Baylor Health Care System (Liver transplant: 002-197, Pregnancy: 009-257, Multiple sclerosis: 009-240, Melanoma: 006-025 & 097-027), Khon Kaen University (Sepsis), the University of Texas Southwestern Medical Center (SoJIA, Dermatomyositis, SLE), Duke University and the Baylor Health Care System (HIV: Duke 8485- 06-4R0 Baylor 006-177), St. Mary’s Hospital London, UK and University of Cape Town, Cape Town, Republic of South Africa (Tuberculosis: St Mary’s REC 06/Q0403/128, U of Cape Town REC 012/2007). The gene expression datasets were selected to cover major classes of immune states (**Table 2**), were required to have at least 25 samples in total, and at least 20% of the total samples were required to be appropriately matched controls.

General descriptions of the study cohorts are as follows: ***S. aureus* cohort:** Children with community-acquired S. aureus infection were enrolled. The clinical syndromes of these patients included skin and soft tissue infection, bacteremia, osteomyelitis, suppurative arthritis, pyomyositis, pneumonia, and disseminated disease. Patients diagnosed with toxic shock syndrome, polymicrobial infections, or treated with corticosteroids in the preceding four weeks were excluded. **Sepsis cohort:** Diagnosis of sepsis was based on accepted international guidelines and defined as presentation with two or more of the following criteria for the systemic inflammatory response syndrome: fever (temperature >38°C or <36°C), tachycardia (heart rate >90 beats/minute), leukocytosis or leukocytopenia (white blood cell count ≥12 × 109/l or ≤4 × 109/l). Blood was collected within 24 hours following the diagnosis of sepsis. Samples were selected for microarray analysis from subjects who had the diagnosis of bacteremic sepsis retrospectively confirmed by the isolation of a causative organism on blood culture. **TB cohort:** Patients were prospectively recruited and sampled, before any anti-mycobacterial treatment was initiated. Active TB disease was confirmed by laboratory isolation of M. tuberculosis on mycobacterial culture of a respiratory specimen (either sputum or bronchoalvelolar lavage fluid). **Influenza cohort:** Children with confirmed influenza infection were recruited. Those with documented bacterial co-infections, children with chronic conditions and systemic steroid treatment within 2 weeks before enrollment were excluded. **RSV cohort:** Patients with confirmed RSV infection were recruited. Children with documented bacterial co-infections, congenital heart disease, chronic lung disease, immunodeficiency, prematurity (<36 wk), systemic steroid treatment within 2 weeks before presentation or additional chronic comorbidities were excluded. **HIV cohort:** Blood samples were obtained from adult patients diagnosed with HIV infection. At enrollment patients were verified as acute Fiebig stages 4 to 6 (plasma RNA+, third generation EIA+, Western blot indeterminant or +). **SLE cohort:** Blood samples were obtained from pediatric patients diagnosed with Systemic Lupus Erythematosus and healthy controls matched for demographic characteristics. MS cohort: Subjects enrolled in the MS cohort had an established diagnosis of relapsing-remitting multiple sclerosis (MS), separately confirmed by an experienced MS neurologist (JTP), exhibited no other health conditions, and had received no treatment(s) for MS, including corticosteroids, for at least 3 months prior to blood draw. **Juvenile Dermatomyositis cohort:** Blood samples were obtained from pediatric patients diagnosed with juvenile dermatomyositis and healthy controls matched for demographic characteristics. **Kawasaki Disease cohort:** Subjects <18 years of age who met the definition of Kawasaki Disease based on the American Heart Association (AHA) criteria (34) were enrolled alongside age and gender matched healthy controls. **Systemic onset Juvenile Arthritis cohort:** Blood samples were obtained from SoJIA patients displaying systemic symptoms only (fever, rash, and/or pericarditis) or displaying systemic symptoms accompanied by arthritis. **COPD cohort:** Enrollment criteria included age over 40 years, no history of concurrent lung cancer, chest surgery, or chronic lung diseases other than COPD (e.g., sarcoidosis, fibrosis, etc.). Participants had no history of allergies or asthma and at the time of initial recruitment had not received oral or systemic corticosteroids during the previous 6 weeks; volunteers were enrolled from three clinics within the Texas Medical Center in Houston, Texas. **B-cell deficiency cohort:** This cohort comprised adults with diagnosis of XLA as documented by markedly reduced numbers of peripheral blood B cells. **Pregnancy cohort:** Pregnant women were recruited at the Baylor Institute for Immunology Research (Dallas, TX) for a study of immunological signatures of pregnancy. **Melanoma cohort:** Enrollment criteria included ages of 21-75 years stage M1a, M1b, M1c biopsy proven metastatic melanoma patients with measurable metastatic lesions by physical exam or scans, acceptable CBC and blood chemistry results, adequate hepatic and renal function, no active CNS metastatic disease. **Liver transplant cohort:** Enrollment criteria included age 17-65 years, having received a liver transplant under maintenance immunosuppression therapy. Subjects in this cohort had not received an acute or chronic rejection diagnosis at the time of sampling.

### RNA extraction and processing

Whole blood for all sample sets were collected into Tempus Blood RNA Tubes (Thermo Fisher Scientific). Total RNA was isolated from whole blood lysate using a MagMAX for Stabilized Blood Tubes RNA Isolation Kit for Tempus Blood RNA Tubes (Thermo Fisher Scientific). RNA quality and quantity were assessed using an Agilent 2100 Bioanalyzer (Agilent Technologies) and a NanoDrop 1000 (NanoDrop Products, Thermo Fisher Scientific). Samples with RNA integrity numbers values >6 were retained for further processing.

### Microarray analysis and data preprocessing

Gene expression profiles from whole blood samples generated using Illumina HumanHT-12 v3.0 or Illumina HumanHT-12 v4.0 expression BeadChips were obtained from 16 groups of patients and controls selected as above. Thus, 16 datasets were used as the input (Table 1). The expression data for each dataset were preprocessed and independently clustered. First, the probes were discarded if they were not present (detection *P* < 0.01) in at least 10 samples or in at least 10% of the samples, whichever was greater. Then, the sample data for each dataset were normalized using the BeadStudio average normalization algorithm. Once normalized, the signal was floored such that all signals <10 were set to 10. Then, the fold change was calculated relative to the median signal for that probe across all samples. If the difference between a signal and the probe’s median signal was <30, or the calculated absolute magnitude of the fold change was <1.2, the fold change was set to 1 to reduce the noise from low-level responses. At this stage, the probes were filtered again. Probes were only retained if they had a calculated absolute fold change >1 in at least 10 samples or in at least 10% of the samples, whichever was greater. Finally, the data was transformed to the log_2_ of the calculated fold changes.

### Module construction algorithm

Sets of coordinately regulated genes, or transcriptional modules, were extracted from the patient’s whole blood microarray datasets. Each of the preprocessed microarray datasets were clustered in parallel using Euclidean distance and the Hartigan’s K-Means clustering algorithm. The ‘ideal’ number of clusters (k) for each dataset was determined within a range of k=1-100 by means of the jump statistic (35). Taking the 16 sets of clusters as the input (**Table 2**), a weighted co-cluster graph was constructed (Chaussabel, Quinn et al. 2008, Chaussabel and Baldwin 2014) To select modules, an iterative algorithm was used to extract the sets of probes that are most frequently clustered together in the same datasets, proceeding from the most stringent requirements to the least, as previously described (Chaussabel, Quinn et al. 2008) This iteration differed from the previous implementation of this algorithm in that the k value was calculated independently for each dataset cluster and the size of the core sub-networks was smaller (10 probes). The algorithm also differed from previous implementations to ensure that the core sub-networks co-clustered in the same datasets. Further details and an example of the code are included in the supplemental methods (**Supplementary File 1**). The resulting 382 module set constitutes the third generation of modular blood transcriptome repertoire constructed since the development of the first generation published in 2008 (4), and the second generation published in 2013 (3).

### Module annotation

Gene Ontology Pathway enrichment:

Module gene lists were investigated using Database for Annotation Visualization and Integrated Discovery (DAVID) version 6.7 (12) This database uses a modified Fisher exact test to identify specific biological/functional categories that are overrepresented in gene sets in comparison with a reference set. The top matched DAVID annotation cluster (using default settings), the top matched canonical pathway from Kyoto encyclopedia of genes and genomes (KEGG), the top matched pathway from BioCarta, and the top matched Gene Ontology biologic process (GO_BP) and molecular function (GO_MF) terms were identified for each module. Each module was also investigated for significant overlap with two other established blood transcriptome module repertoires (3,9). The findings are summarized in the module annotation spreadsheet (**Supplemental File 2**).

Gene Set Annotation (GSAn):

To further annotate the modules, a new alternative to statistical enrichment analysis tool called GSAn was applied (13). Statistical enrichment methods may have limitations (36–38), as these methods tend to focus on the subpart of the most studied genes and to provide gene set annotation results that cover a limited number of the well-annotated genes. To address these issues, GSAn offers: (i) an original method that combines semantic similarity measures and data mining approaches to perform a unified and synthetic annotation for a gene set of interest, and (ii) a visualization approach to facilitate the interactive exploration of the gene set annotation results according to the hierarchical structure of Gene Ontology (13). The tool is available online: https://gsan.labri.fr/. A page listing analysis results for all 382 generation 3 blood transcriptome modules can be accessed here: https://ayllonbe.github.io/modulesV3/index.html.

Pathway enrichment analyses:

These analyses were conducted using Ingenuity Pathway Analysis (Qiagen, Valencia, CA).

Literature profiling:

Literature Lab(tm) (LitLab; from Acumenta Biotech, Boston, MA) was used to associate genes within a particular module to terms used in PubMed abstracts (Febbo, Mulligan et al. 2007). Association scores reflecting the strength of the associations were used to calculate the “Product Scores”. The top three terms that showed the strongest association and highest “Product Scores” were used to create the functional annotation. A similar approach using LitLab has been previously reported (Alsina, Israelsson et al. 2014). The steps taken to annotate all 382 modules is described here in brief. All statistical analyses were performed using Microsoft Excel (2010) with Visual Basic for Applications (VBA), Linux-based command line in Mac OS, and R statistical software.

To construct a Product Scores Table, all the terms available in LitLab (>80,000) were listed. Then, the genes in each module were submitted as a list to LitLab Editor and manually validated using LitLab’s built-in validation tool and/or NCBI Gene (https://www.ncbi.nlm.nih.gov/gene) prior to submission for analysis using all domains available. After completing the analysis, the summary result page was exported to an xls file. Using the UNIX command line, the exported files were converted to csv files with the filename appended in the last column of each row and vertically appended. The “merged” file was used to populate the table that included all the available LitLab terms. The top three terms with the highest Product Scores were selected to represent the module functional annotation and are tabulated in column I of the module annotation table (**Supplementary file 2**).

### Fingerprint grid plot visualization

Modules were arranged on a grid based on their similarities in patterns of activity across the 16 input datasets, each of them corresponding to a different pathological or physiological state. First, the modules were partitioned using k-means clustering, which resulted in the constitution of 38 clusters. Given the possibility of collapsing values of the modules constituting each cluster in a single “aggregate” value, the term “module aggregate” was used to designate each cluster (A1 to A38). Of these 38 k-means clusters, 27 comprised >1 module. The modules were then arranged on a grid with each row corresponding to modules belonging to the same aggregate (Figure 4). The total number of rows on the grid equaled 27 and number of columns equaled the largest number of modules for a given aggregate (42 for aggregate A2). For each module, the highest of the two values indicating an increase or a decrease was selected for visualization (e.g. if % increase > % decrease, then a red sport representing % increase is shown).

### Module Fingerprint analysis and visualization

The modular analysis was performed by using 14,168 transcripts. Fold change was computed using gene expression data prior log2 transformation. For group comparisons, paired t-test was performed on log2-transformed data (Fold change (FC) cut off = 1.5; FDR cut off = 0.1). For individual-patients analysis, each sample was compared to mean of control samples in each dataset. Cut off comprised a absolute FC > 1.5 and a difference in gene expression level > 10. A module was considered to be “responsive” when the proportion of differentially expressed transcripts (as defined above) was greater than 15%. Data visualization were performed using “ComplexHeatmap” (36).

### Generation of Circular Plots

Circular plots were generated to represent module expression at the gene-level (**Figure 7**). Fold change in case vs control group in each dataset were shown. Probes that passed FDR < 0.1 and FC > 1.5 were present in gradated red colors and those with FC < -1.5 were represented in gradated blue colors. The position of the genes on each of the circular plots is fixed.

### Statistical analyses

Numerical data were processed and analyzed using R statistical software (version1.1.463-©2009- 2018 RStudio, Inc). Student’s t-test was used to determine which group was different. Values of *p < 0*.*05* were considered to indicate statistical significance, with adjustment by multiple-testing correction when needed (FDR, Benjamini–Hochberg procedure).

### Availability of data and material

Transcriptome profiling data are deposited in the Gene Expression Omnibus, https://www.ncbi.nlm.nih.gov/geo/, under the accession GSE100150.

## SUPPLEMENTAL INFORMATION

**Supplementary File 1: Module generation method and pseudocode**

This Word document describes the algorithm used to construct this modular repertoire framework along with pseudocode, which may be used as a basis for implementation in programming languages such as R or Python.

**Supplementary File 2: Annotated module repertoire**

This Excel spreadsheet lists the 382 modules constituting this third generation blood transcriptome module repertoire. Included are the number of genes, the list of member genes by symbol and the probe ID, summarized functional annotations.

**Supplementary File 3: Module fingerprints of all 16 pathological and physiological states**

This PDF file contains the modular fingerprints generated for each of the 16 input datasets.

