## Supplementary File 1 for "Development and Characterization of a Fixed Repertoire of Blood Transcriptome Modules Based on Co-expression Patterns Across Immunological States"

### **Supplemental File 1: Module generation and pseudocode**

Each of the preprocessed microarray datasets was clustered in parallel using Euclidean distance and the Hartigan’s K-Means clustering algorithm, a hybrid of hierarchical and k-means clustering algorithms. An additional modification was made to the algorithm. If at any k the algorithm creates a cluster whose members’ average Pearson correlation to the mean cluster vector is <0.3, the cluster is deleted and the algorithm begins again at k-1. The ‘ideal’ number of clusters (k) for each dataset was determined within a range of k=1-100 by means of the jump statistic.

Taking the 16 sets of clusters as the input, a weighted co-cluster graph — a probe by probe matrix where the value of each cell is set to a 16-bit bitmask (each bit corresponding to a dataset) indicating in which datasets the probes were co-clustered — was created. The bit was set to 1 if the probe pair co-clusters in the corresponding dataset, and to 0 if not. Therefore, the number of times the probes co-clustered (the weight) is equivalent to the number of bits in the mask that are set to 1. At this point, the goal is to extract sets of probes that are most frequently clustered together in the same datasets, proceeding from the most stringent requirements to the least. To accomplish this, an iterative algorithm was used. To begin, the maximum clique threshold was initialized to the number of input cluster sets, the paraclique threshold (pt) was calculated, and a minimum seed size was chosen (we used fifteen). The outer loop began by creating an unweighted graph by applying the maximum clique threshold (mct) to the weighted co-cluster graph such that a probe pair, or edge, was connected in the unweighted graph only if the corresponding weight in the co-cluster graph equaled or exceeded this threshold.

For the inner loop, the first step was to isolate the largest set of probes such that all probes in the set were completely connected in the unweighted graph. In graph theoretical terms, the probes form a maximum clique. An additional constraint was imposed that all probes in the set must co-cluster in the same datasets at least mct times by taking the intersection of the bitmasks for every edge in the clique and the result must contain at least mct bits set to 1. This new bitmask was the common co-cluster bitmask for the clique.

If the size of the clique was smaller than the minimum seed size, the inner loop was escaped, reduced mct by one, and the process returned to the beginning of the outer loop; otherwise, it became the seed for a module. To allow for inevitable clustering inaccuracies, the paraclique algorithm was used. The co-cluster graph was re-visited and added to the seed any probe that was found to co-cluster with at least 90% of the seed’s members in at least pt of the datasets represented in the clique’s common co-cluster bitmask. This final probe set was a module: it was removed from both graphs and named in accordance with the iterations in which it was found (i.e. a module extracted in the first iteration of the outer loop and the second iteration of the inner loop is designated M1.2). The inner loop then began again with the reduced graphs. The Pseudocode is shown below:

Integer nLastQuartile = 4;

Integer nMaxRelaxtion = m_nNumDatasets / 3;

Integer nRelaxtionIncrement = Math.max(1, (nMaxRelaxtion / 3));

Integer nRelaxtion = nMaxRelaxtion;

for (int nCliqueThreshold = numberOfDatasets; nCliqueThreshold >= 1; nCliqueThreshold--)

{

Integer nQuartile = ((nThreshold * 100) / m_nNumDatasets) / 25;

if (nQuartile.equals(nLastQuartile) == false)

{

if (nQuartile <= 2)

{

nRelaxtion = Math.max(0, nRelaxtion - nRelaxtionIncrement);

}

nLastQuartile = nQuartile;

}

Integer nParacliqueThreshold = nThreshold - nRelaxtion;

do

{

maximumClique = find maximum clique w co-clustering weight >= nCliqueThreshold

if (size of maximumClique > 15)

{

paraclique = find paraclique in graph

remove maximumClique and paraclique from graph

}

}	while (maximumClique is found)
