## Supplementary File 3 for "Development and Characterization of a Fixed Repertoire of Blood Transcriptome Modules Based on Co-expression Patterns Across Immunological States"

GSE100156: Bcell vs Control

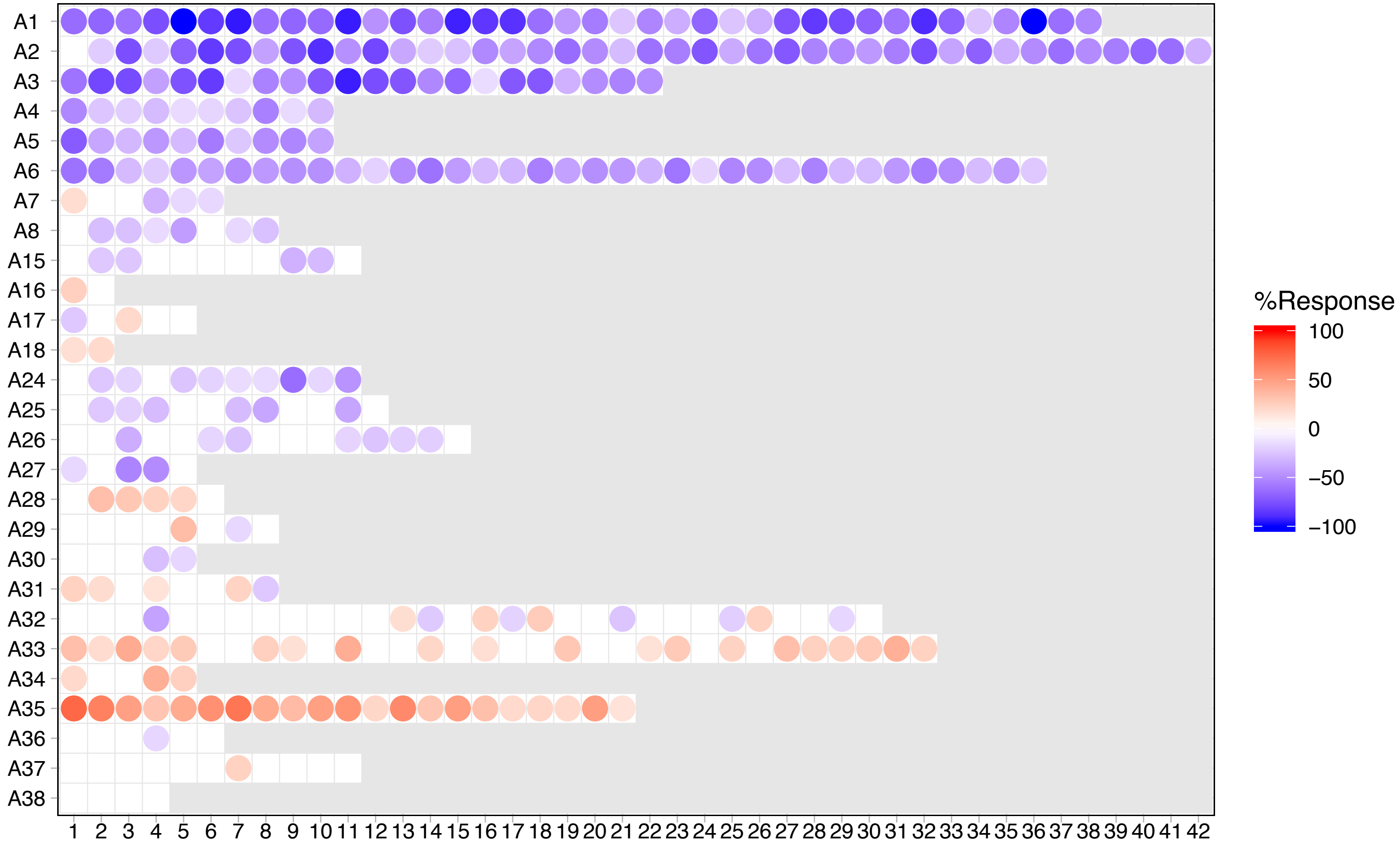

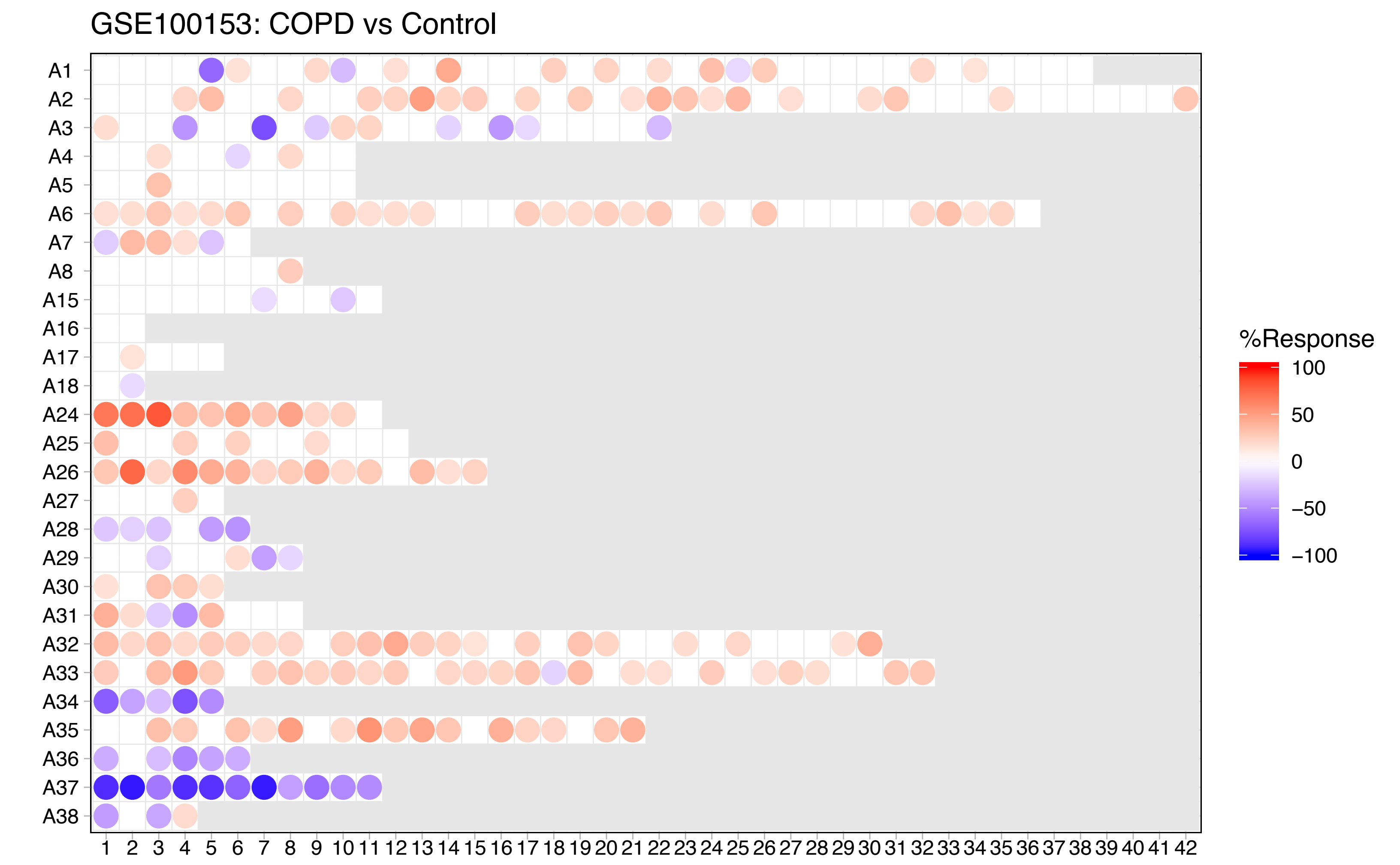
GSE100153: COPD vs Control
A1
A2
A3
A4
A5
A6
A7
A8
A15
A16
A17
A18
A24
A25
A26
A27
A28
A29
A30
A31
A32
A33
A34
A35
A36
A37
A38
1 2 3 4 5 6 7 8 9 10 11 12 13 14 15 16 17 18 19 20 21 22 23 24 25 26 27 28 29 30 31 32 33 34 35 36 37 38 39 40 41 42
%Response
100
50
0
−50
−100

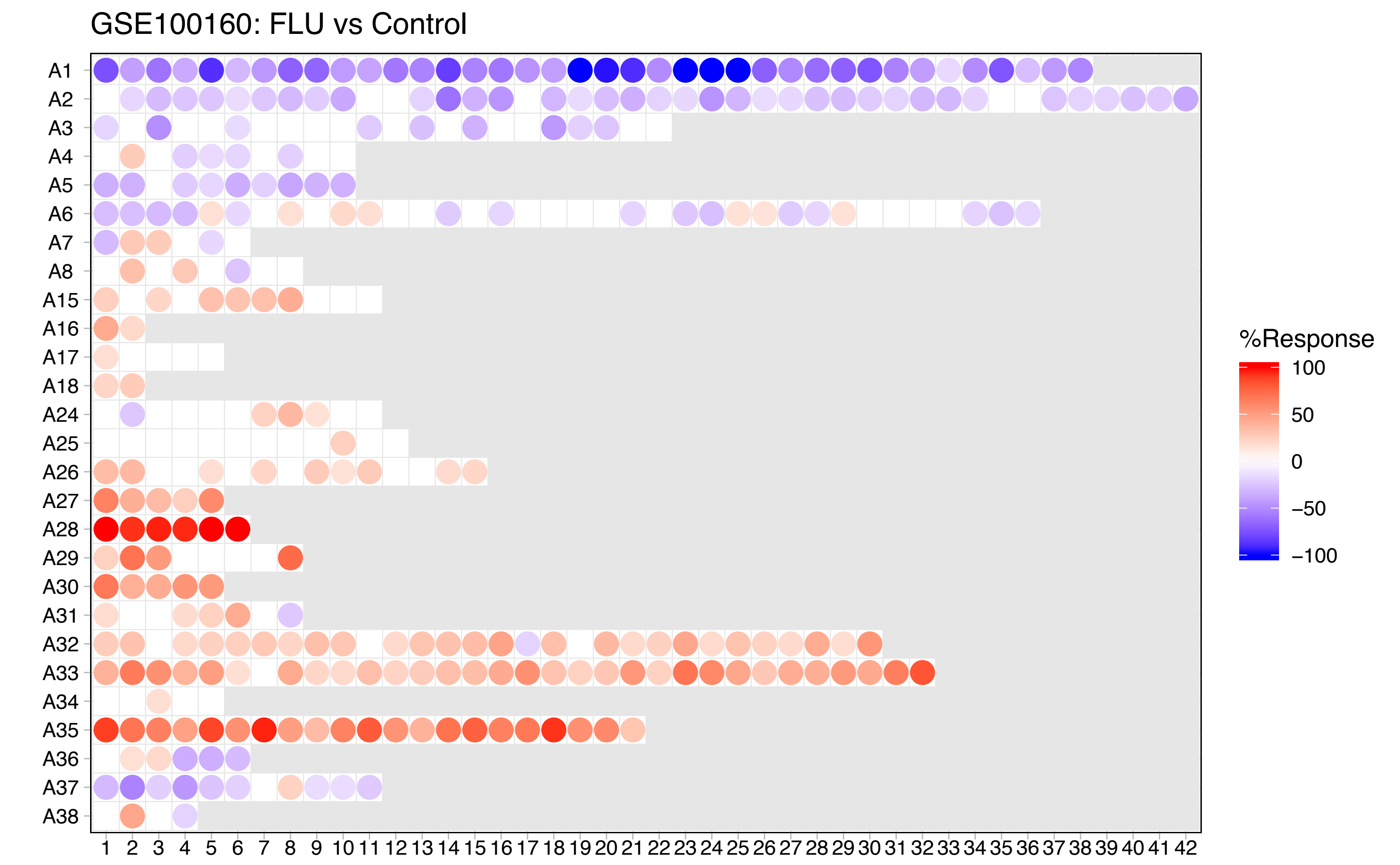
GSE100160: FLU vs Control
A1
A2
A3
A4
A5
A6
A7
A8
A15
A16
A17
A18
A24
A25
A26
A27
A28
A29
A30
A31
A32
A33
A34
A35
A36
A37
A38
1 2 3 4 5 6 7 8 9 10 11 12 13 14 15 16 17 18 19 20 21 22 23 24 25 26 27 28 29 30 31 32 33 34 35 36 37 38 39 40 41 42
%Response
100
50
0
−50
−100

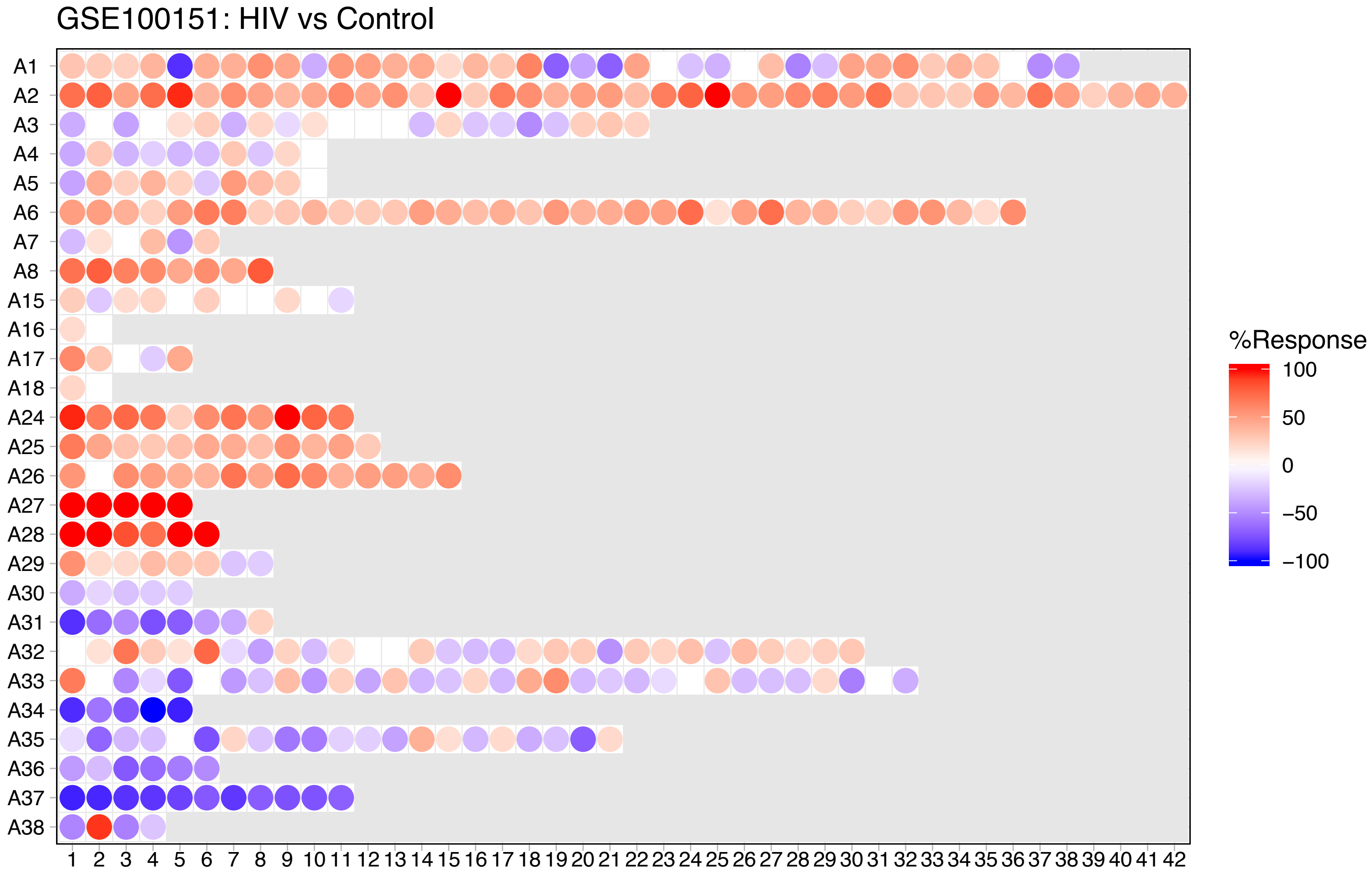

GSE100151: HIV vs Control
A1
A2
A3
A4
A5
A6
A7
A8
A15
A16
A17
A18
A24
A25
A26
A27
A28
A29
A30
A31
A32
A33
A34
A35
A36
A37
A38
1 2 3 4 5 6 7 8 9 10 11 12 13 14 15 16 17 18 19 20 21 22 23 24 25 26 27 28 29 30 31 32 33 34 35 36 37 38 39 40 41 42
%Response
100
50
0
−50
−100

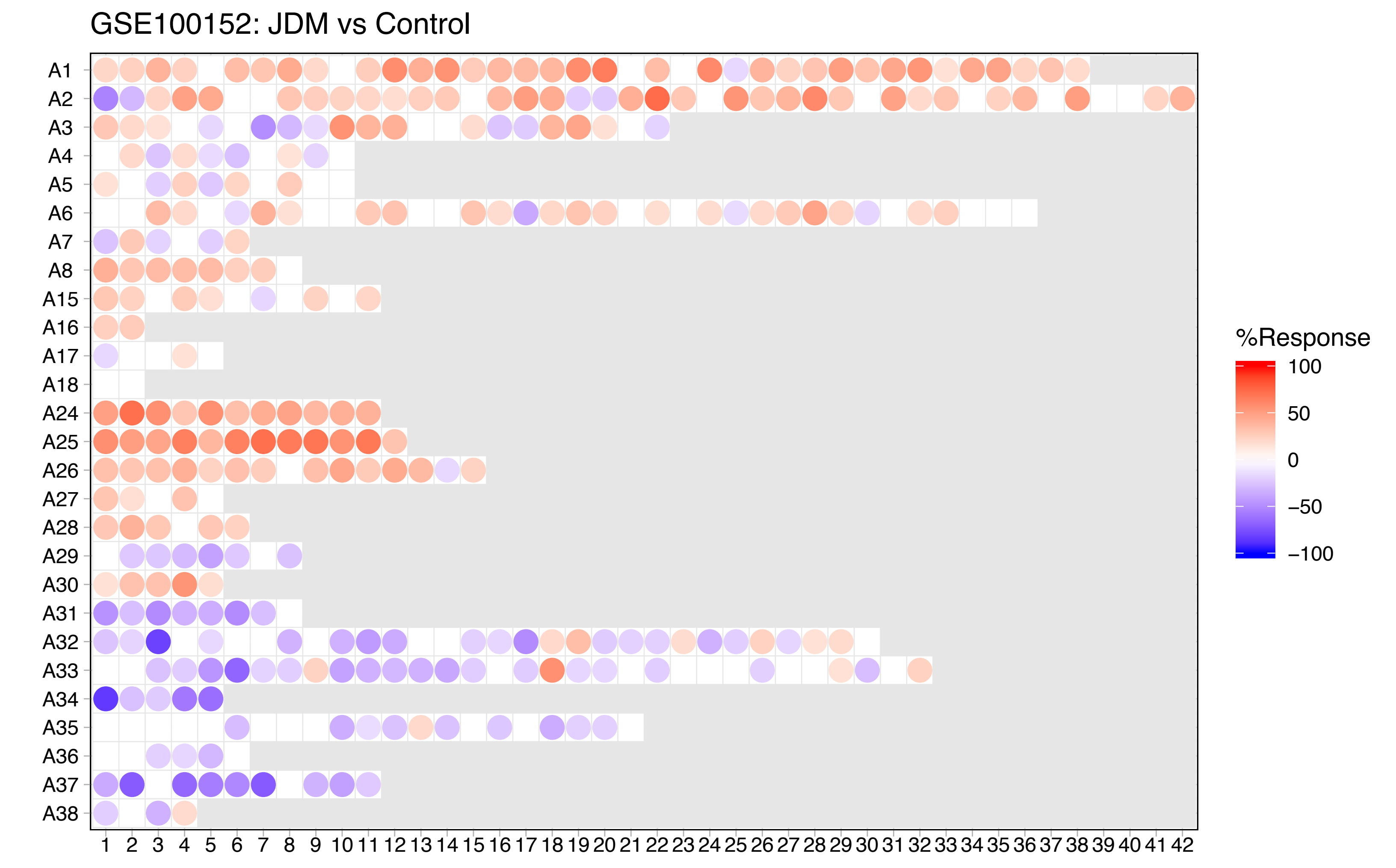
GSE100152: JDM vs Control
A1
A2
A3
A4
A5
A6
A7
A8
A15
A16
A17
A18
A24
A25
A26
A27
A28
A29
A30
A31
A32
A33
A34
A35
A36
A37
A38
1 2 3 4 5 6 7 8 9 10 11 12 13 14 15 16 17 18 19 20 21 22 23 24 25 26 27 28 29 30 31 32 33 34 35 36 37 38 39 40 41 42
%Response
100
50
0
−50
−100

GSE100154: Kawasaki vs Control

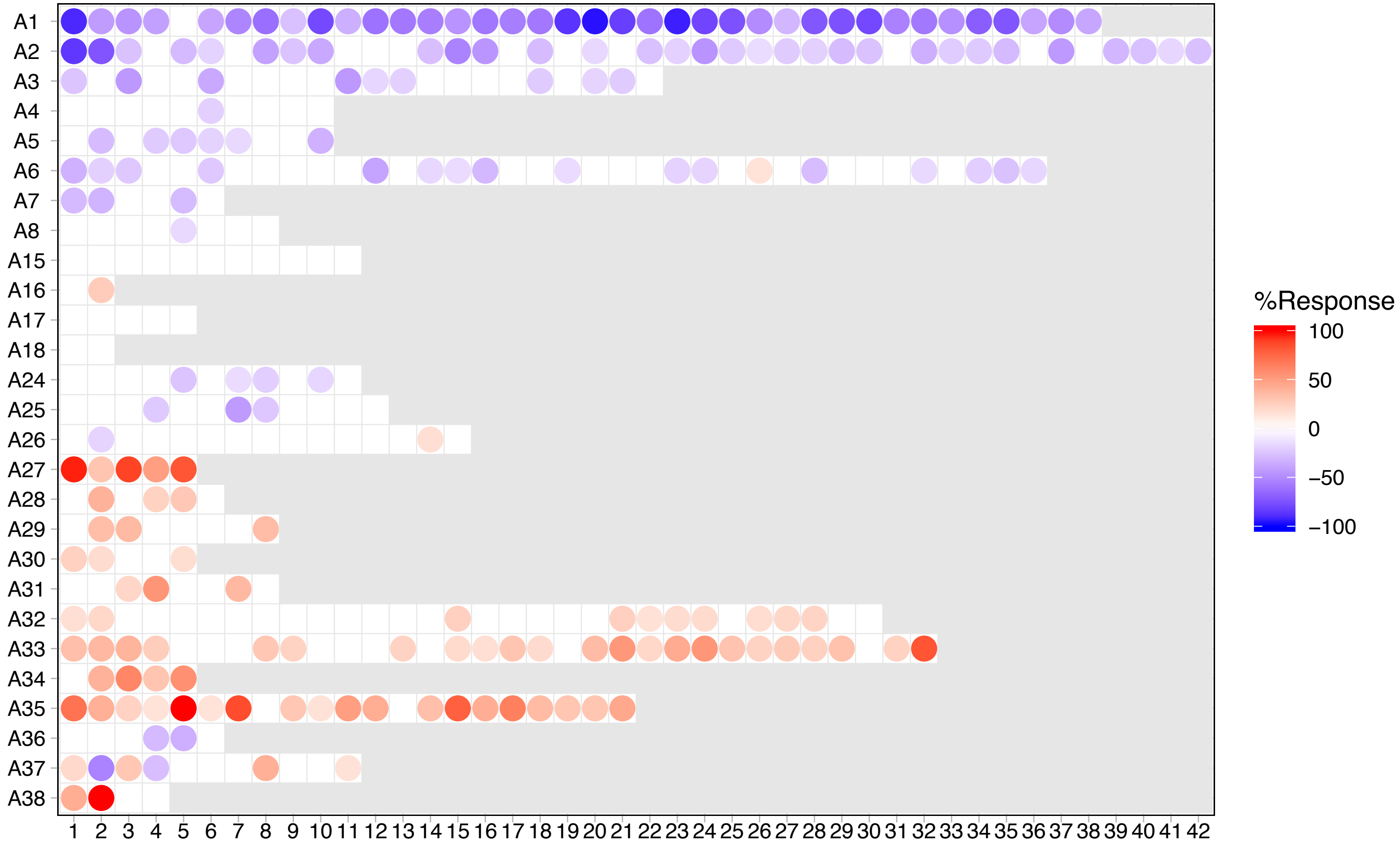

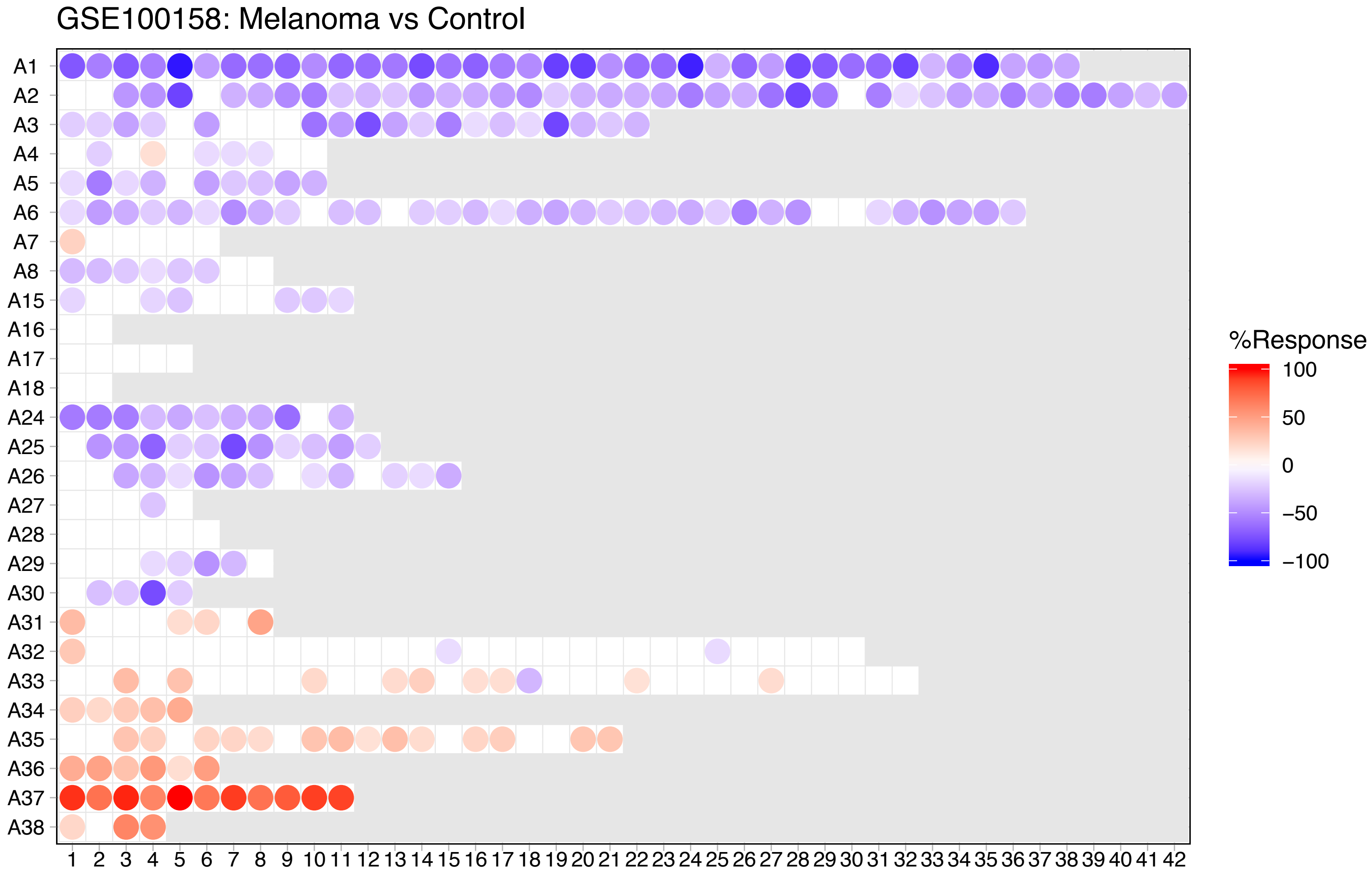
GSE100158: Melanoma vs Control
A1
A2
A3
A4
A5
A6
A7
A8
A15
A16
A17
A18
A24
A25
A26
A27
A28
A29
A30
A31
A32
A33
A34
A35
A36
A37
A38
1 2 3 4 5 6 7 8 9 10 11 12 13 14 15 16 17 18 19 20 21 22 23 24 25 26 27 28 29 30 31 32 33 34 35 36 37 38 39 40 41 42
%Response
100
50
0
−50
−100

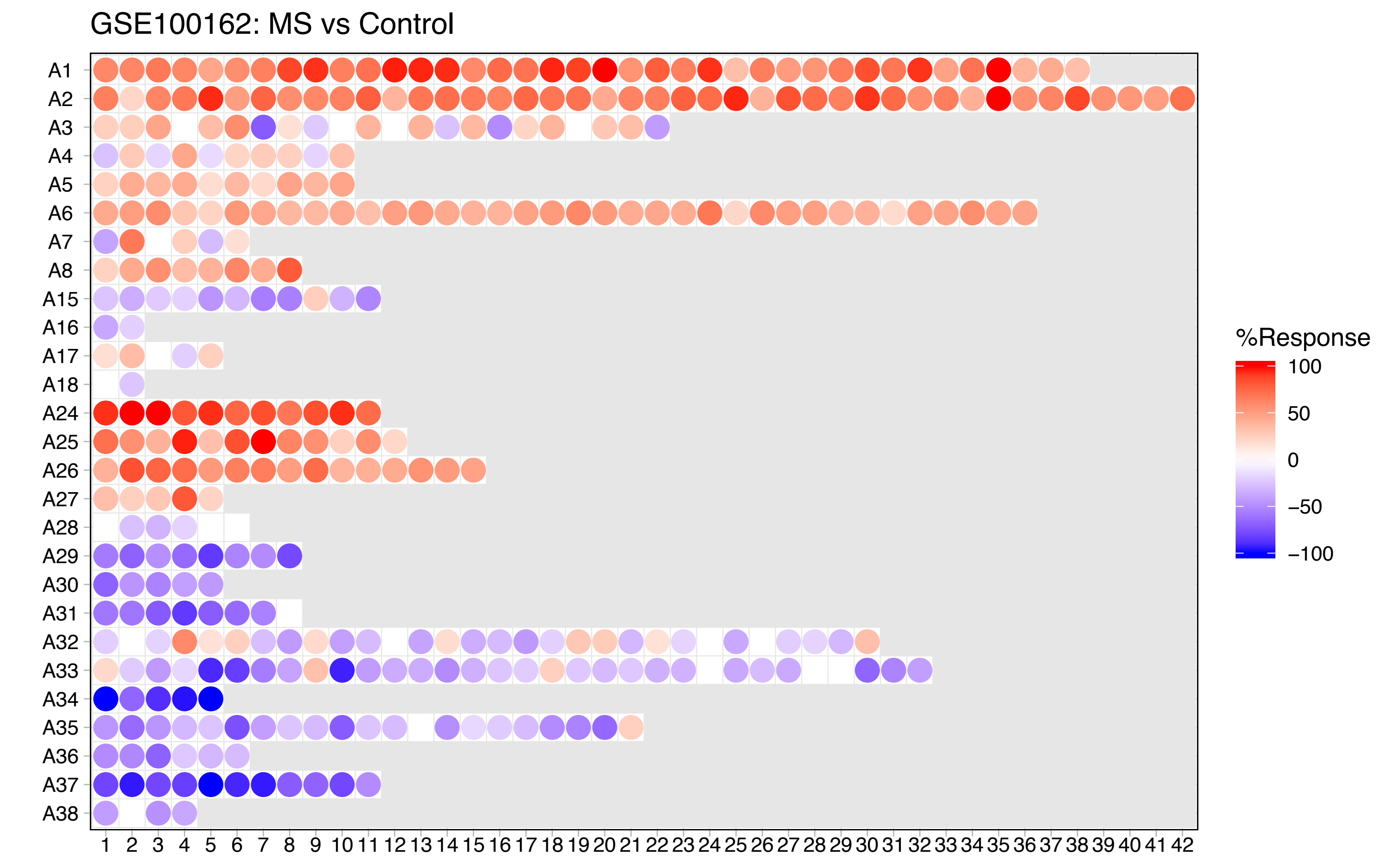

GSE100162: MS vs Control
A1
A2
A3
A4
A5
A6
A7
A8
A15
A16
A17
A18
A24
A25
A26
A27
A28
A29
A30
A31
A32
A33
A34
A35
A36
A37
A38
1 2 3 4 5 6 7 8 9 10 11 12 13 14 15 16 17 18 19 20 21 22 23 24 25 26 27 28 29 30 31 32 33 34 35 36 37 38 39 40 41 42
%Response
100
50
0
−50
−100

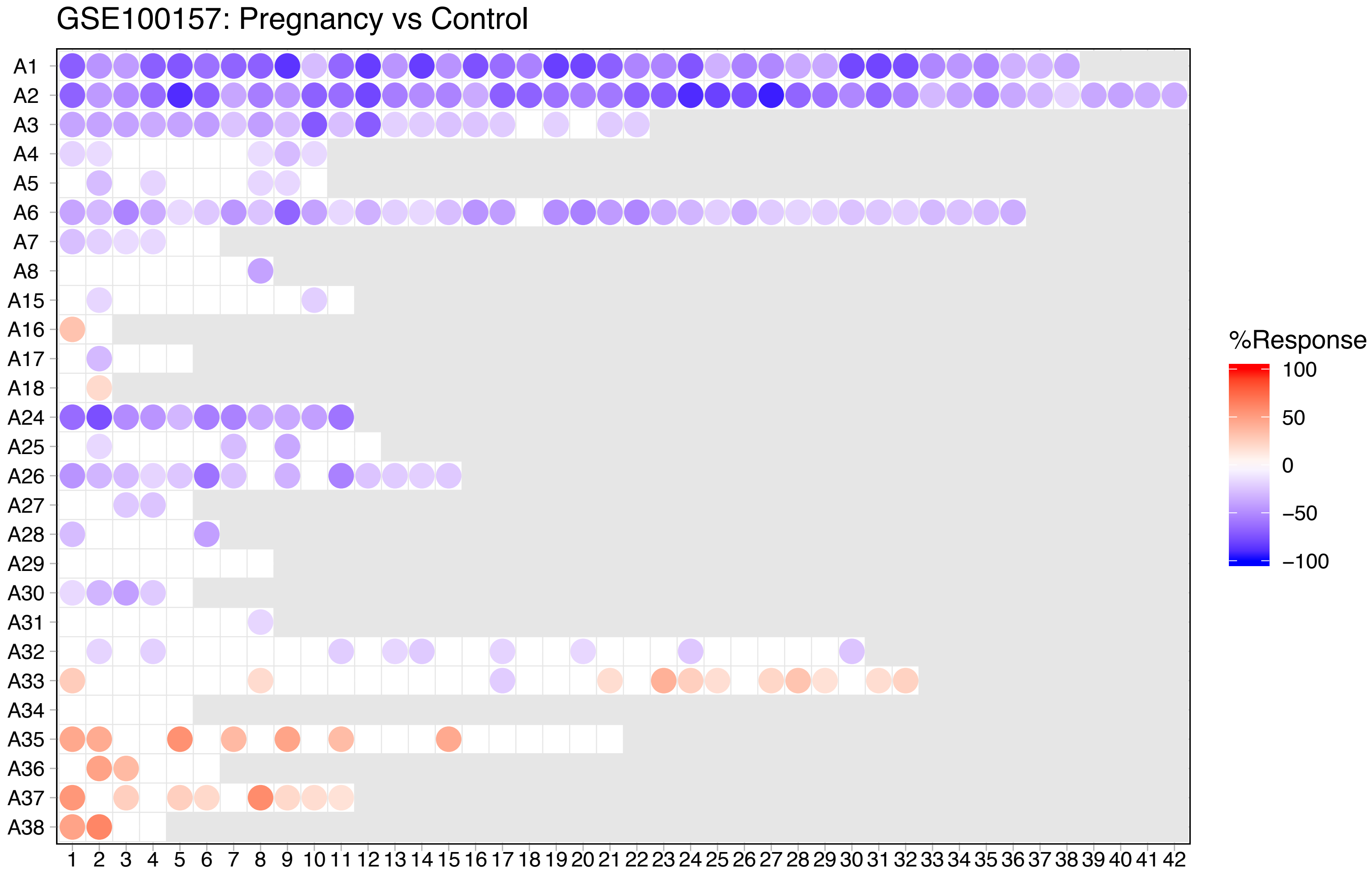
GSE100157: Pregnancy vs Control
A1
A2
A3
A4
A5
A6
A7
A8
A15
A16
A17
A18
A24
A25
A26
A27
A28
A29
A30
A31
A32
A33
A34
A35
A36
A37
A38
1 2 3 4 5 6 7 8 9 10 11 12 13 14 15 16 17 18 19 20 21 22 23 24 25 26 27 28 29 30 31 32 33 34 35 36 37 38 39 40 41 42
%Response
100
50
0
−50
−100

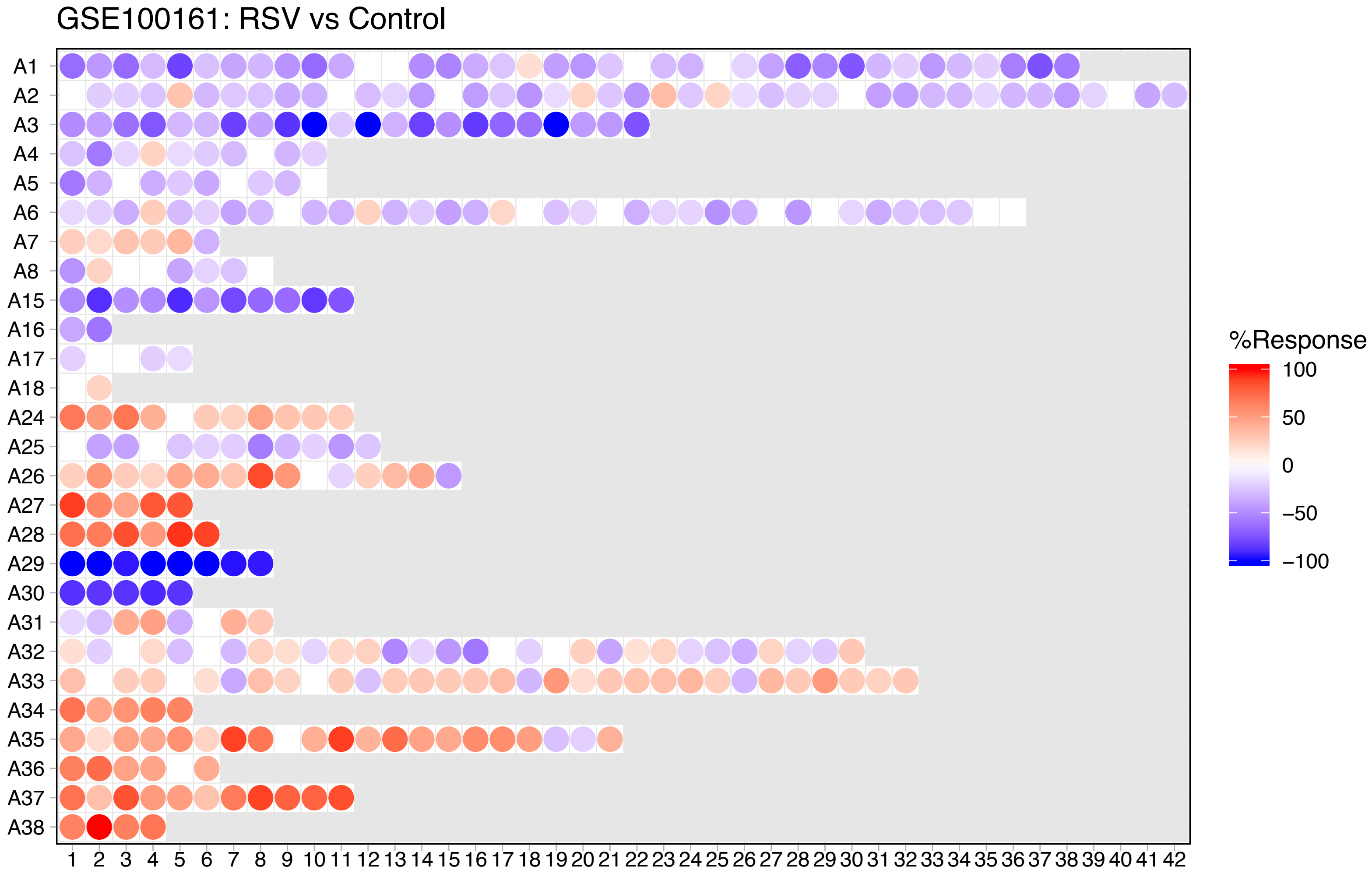
GSE100161: RSV vs Control
A1
A2
A3
A4
A5
A6
A7
A8
A15
A16
A17
A18
A24
A25
A26
A27
A28
A29
A30
A31
A32
A33
A34
A35
A36
A37
A38
1 2 3 4 5 6 7 8 9 10 11 12 13 14 15 16 17 18 19 20 21 22 23 24 25 26 27 28 29 30 31 32 33 34 35 36 37 38 39 40 41 42
%Response
100
50
0
−50
−100

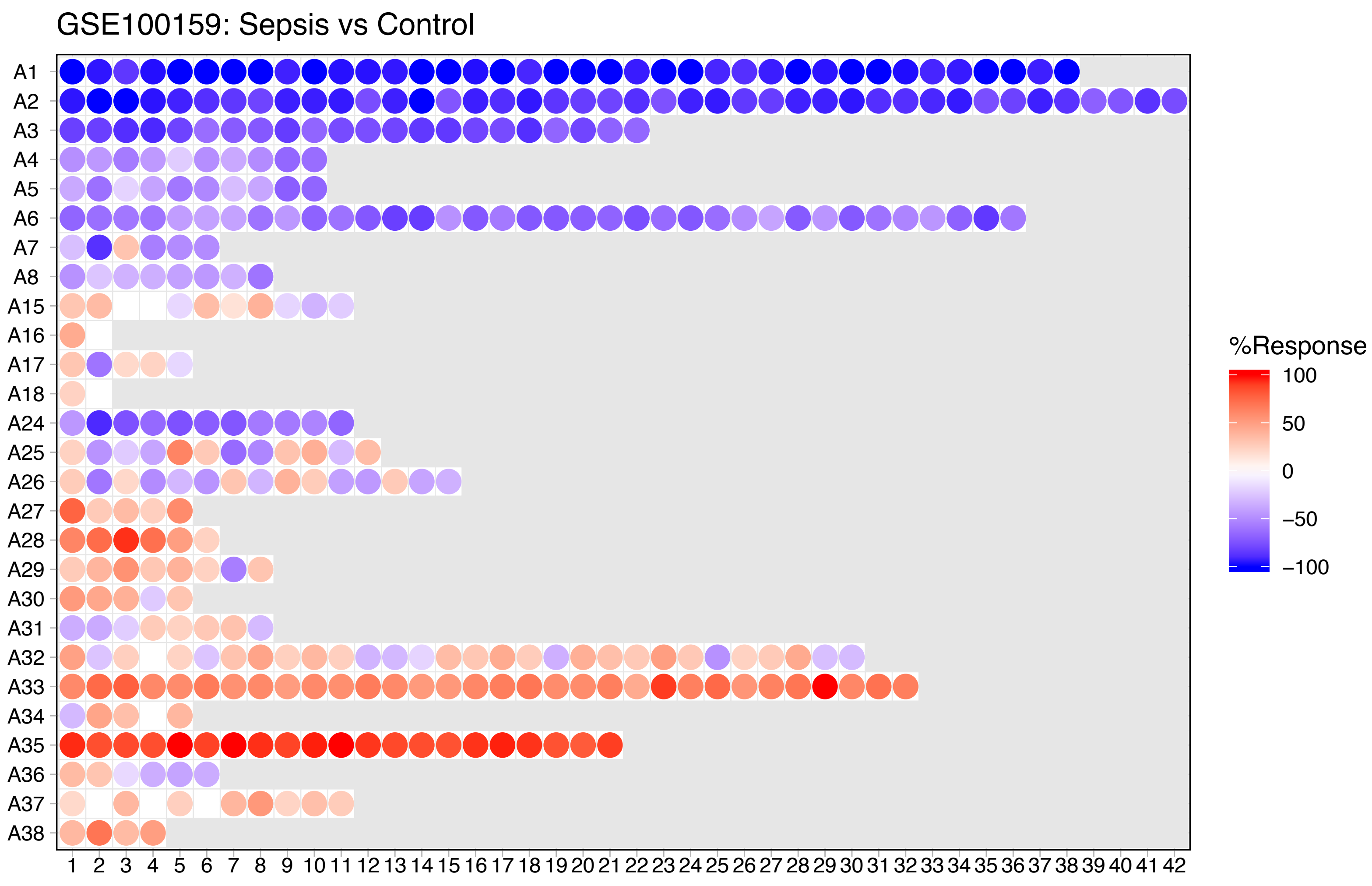
GSE100159: Sepsis vs Control
A1
A2
A3
A4
A5
A6
A7
A8
A15
A16
A17
A18
A24
A25
A26
A27
A28
A29
A30
A31
A32
A33
A34
A35
A36
A37
A38
1 2 3 4 5 6 7 8 9 10 11 12 13 14 15 16 17 18 19 20 21 22 23 24 25 26 27 28 29 30 31 32 33 34 35 36 37 38 39 40 41 42
%Response
100
50
0
−50
−100

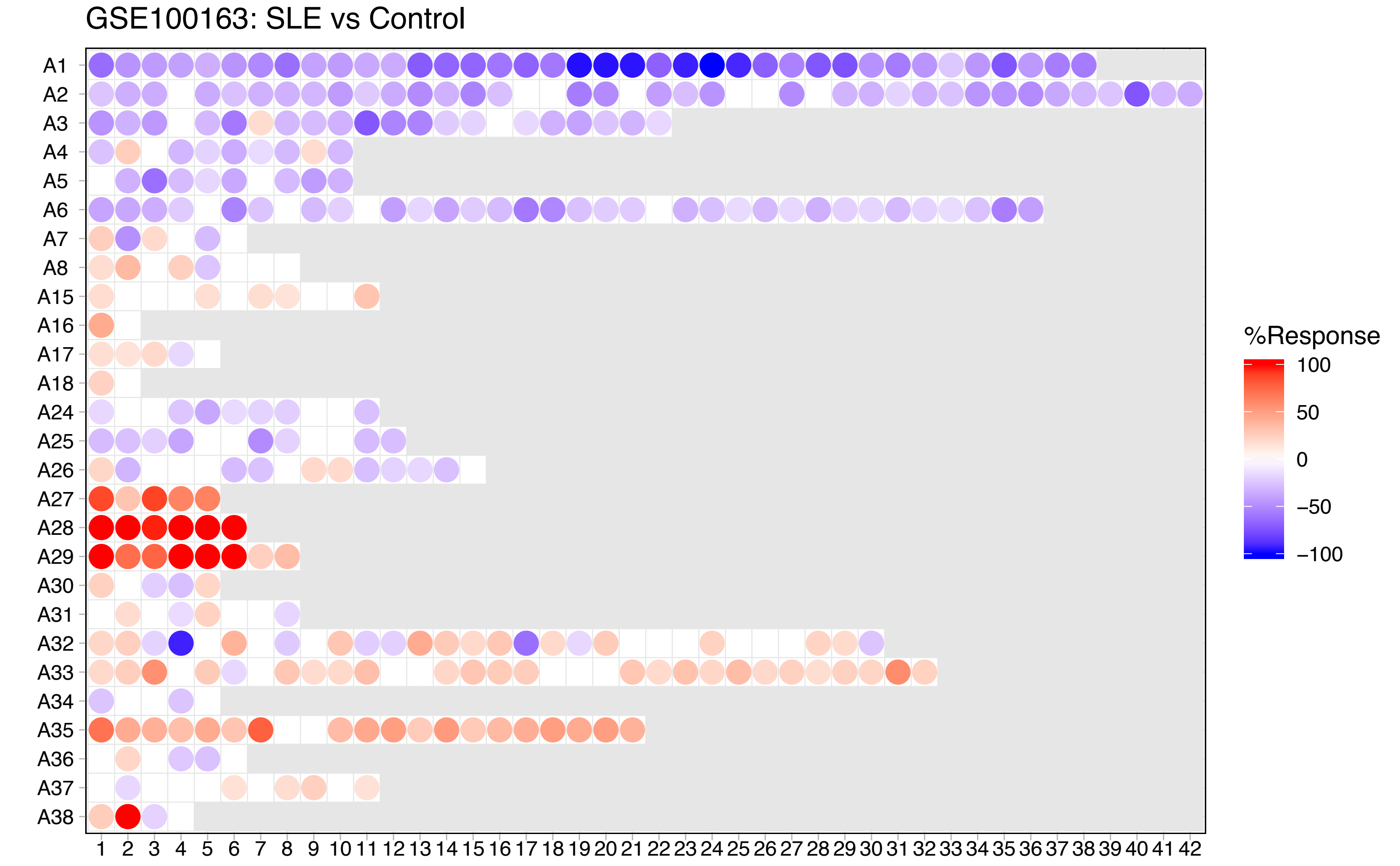

GSE100163: SLE vs Control
A1
A2
A3
A4
A5
A6
A7
A8
A15
A16
A17
A18
A24
A25
A26
A27
A28
A29
A30
A31
A32
A33
A34
A35
A36
A37
A38
1 2 3 4 5 6 7 8 9 10 11 12 13 14 15 16 17 18 19 20 21 22 23 24 25 26 27 28 29 30 31 32 33 34 35 36 37 38 39 40 41 42
%Response
100
50
0
−50
−100

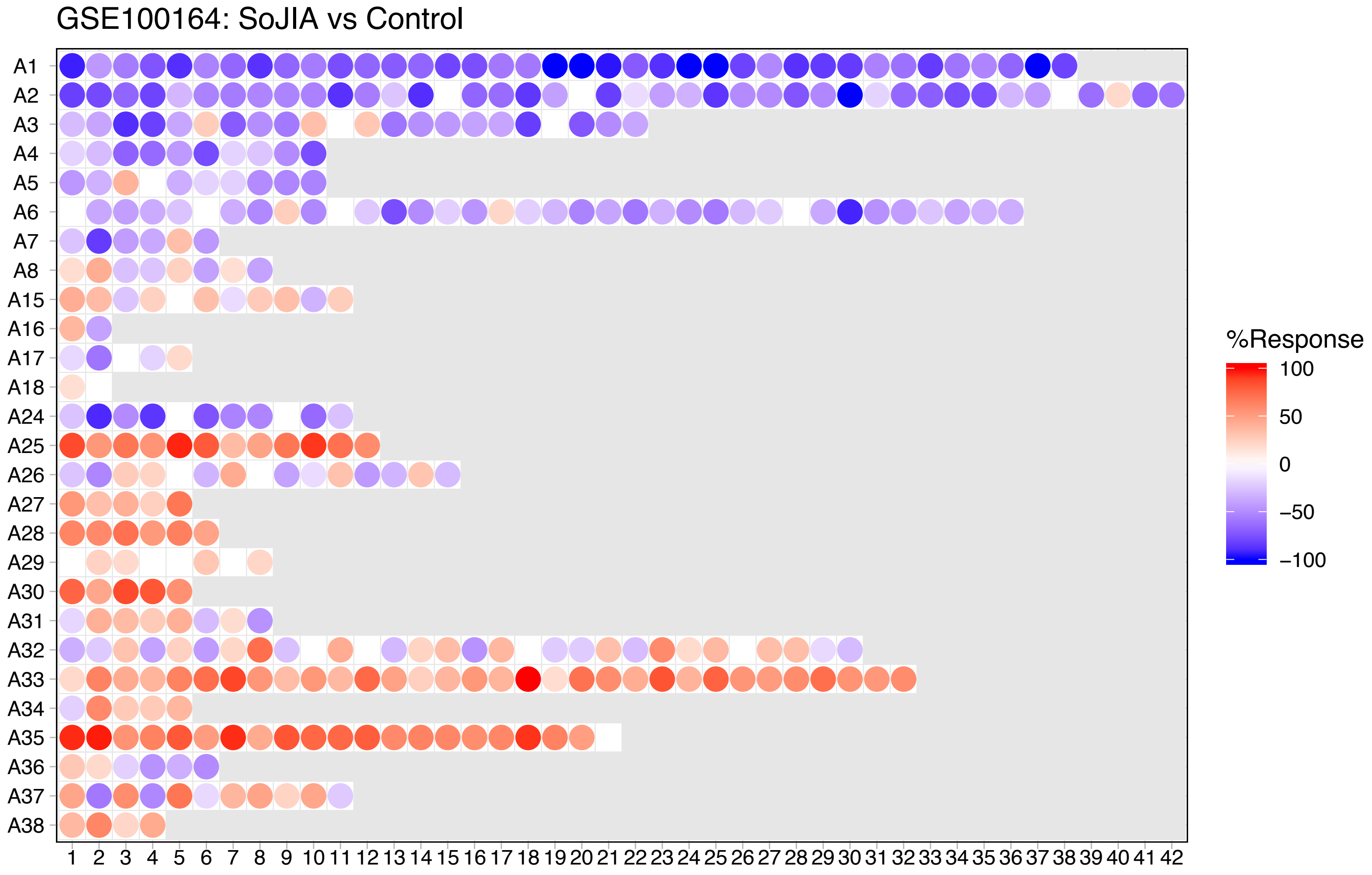

GSE100164: SoJIA vs Control
A1
A2
A3
A4
A5
A6
A7
A8
A15
A16
A17
A18
A24
A25
A26
A27
A28
A29
A30
A31
A32
A33
A34
A35
A36
A37
A38
1 2 3 4 5 6 7 8 9 10 11 12 13 14 15 16 17 18 19 20 21 22 23 24 25 26 27 28 29 30 31 32 33 34 35 36 37 38 39 40 41 42
%Response
100
50
0
−50
−100

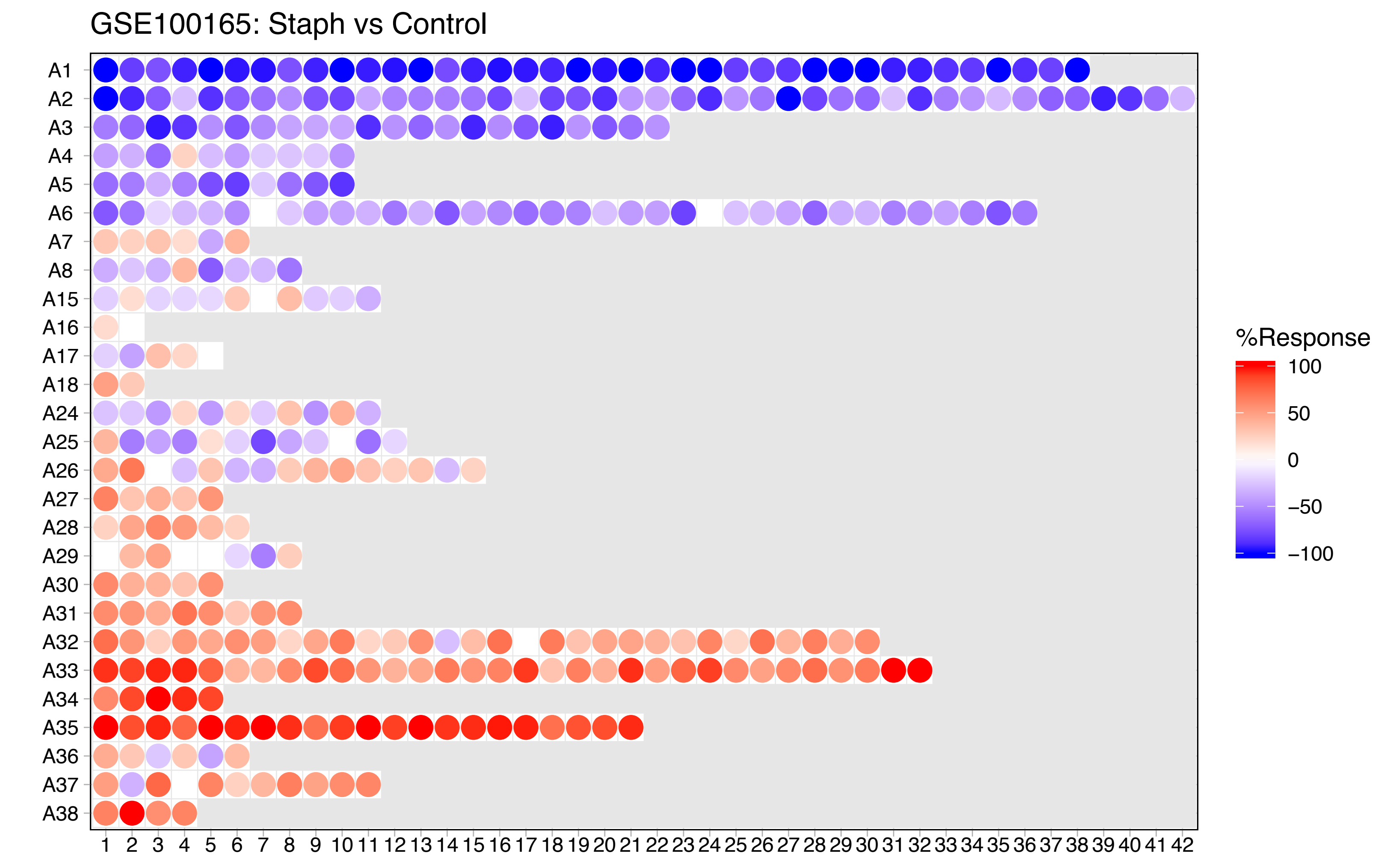

GSE100165: Staph vs Control
A1
A2
A3
A4
A5
A6
A7
A8
A15
A16
A17
A18
A24
A25
A26
A27
A28
A29
A30
A31
A32
A33
A34
A35
A36
A37
A38
1 2 3 4 5 6 7 8 9 10 11 12 13 14 15 16 17 18 19 20 21 22 23 24 25 26 27 28 29 30 31 32 33 34 35 36 37 38 39 40 41 42
%Response
100
50
0
−50
−100

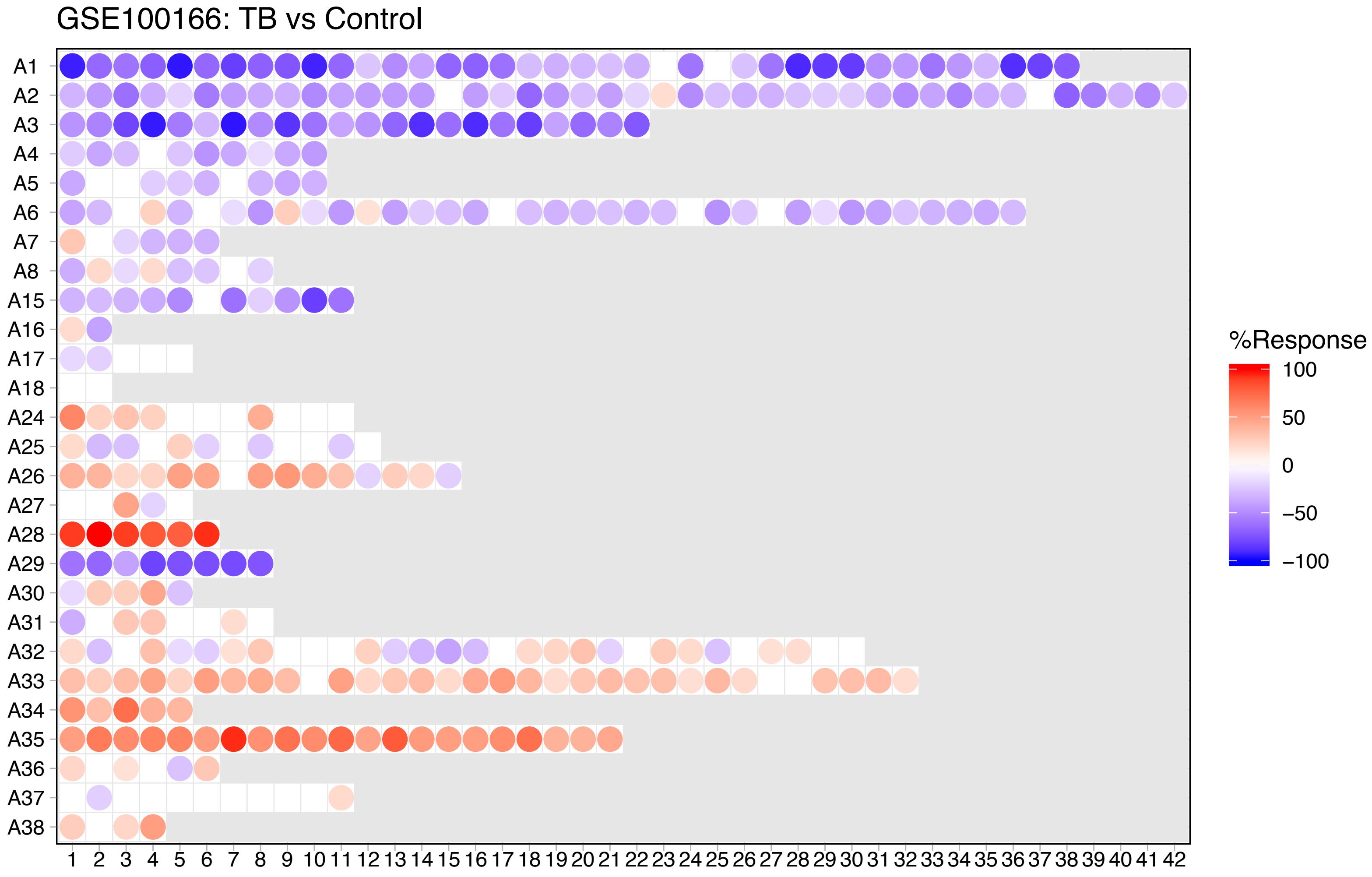
GSE100166: TB vs Control
A1
A2
A3
A4
A5
A6
A7
A8
A15
A16
A17
A18
A24
A25
A26
A27
A28
A29
A30
A31
A32
A33
A34
A35
A36
A37
A38
1 2 3 4 5 6 7 8 9 10 11 12 13 14 15 16 17 18 19 20 21 22 23 24 25 26 27 28 29 30 31 32 33 34 35 36 37 38 39 40 41 42
%Response
100
50
0
−50
−100

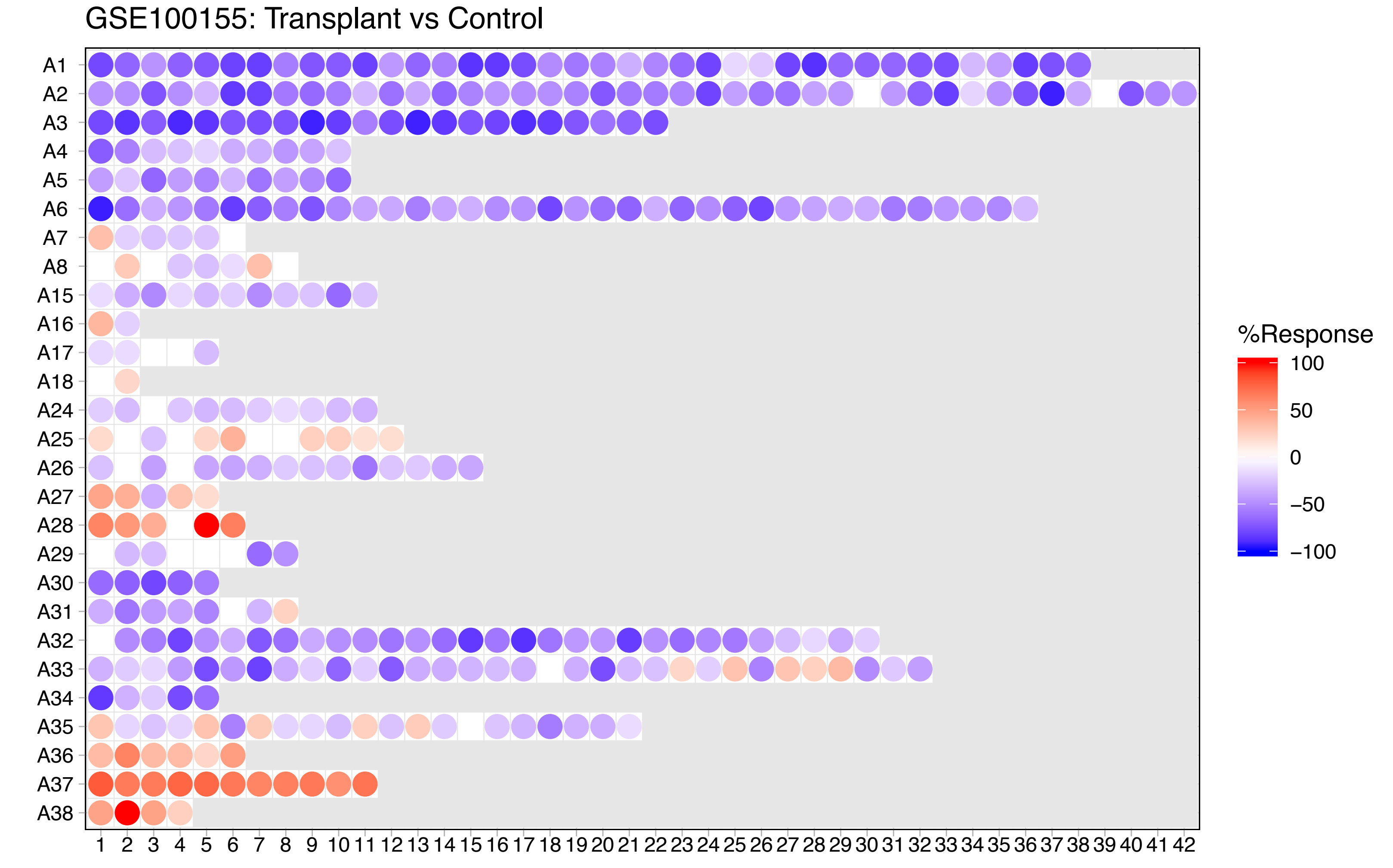

GSE100155: Transplant vs Control
A1
A2
A3
A4
A5
A6
A7
A8
A15
A16
A17
A18
A24
A25
A26
A27
A28
A29
A30
A31
A32
A33
A34
A35
A36
A37
A38
1 2 3 4 5 6 7 8 9 10 11 12 13 14 15 16 17 18 19 20 21 22 23 24 25 26 27 28 29 30 31 32 33 34 35 36 37 38 39 40 41 42
%Response
100
50
0
−50
−100
